# Peptide-functionalized fluorescent polymeric nanoparticles: polysarcosine length defines stealth properties

**DOI:** 10.1101/2025.04.20.649688

**Authors:** Tanushree Gupta, Quynh Anh Bui, Horgan Manirakiza, Lina El Hajji, Nicolas Humbert, Abdel Wahab Mouhamad, Andreas Reisch, Arnaud Gautier, Andrey S. Klymchenko

## Abstract

Stealth properties of nanoparticles are essential for their proper functionalization in biological systems. To address limitations of polyethylene glycol (PEG), commonly used for this purpose, we explore the potential of polysarcosine (PSar) as stealth shell in peptide-functionalized dye-loaded polymeric NPs. To this end, polymeric NPs loaded with rhodamine dye with bulky hydrophobic counterion and bearing azide groups at their surface were grafted with PSar of different lengths ranging from 5 to 19 sarcosine units using strain-promoted cycloaddition. The obtained peptide-functionalized NPs showed remarkable colloidal stability in physiological media. The length of PSar showed a profound effect on stealth properties of NPs. The increase in the length of grafted PSar lead to decrease in the negative surface charge to nearly neutral values and decreased protein adsorption according to fluorescence correlation spectroscopy. The NPs with 19mer PSar showed minimal interactions with live cells and glass surfaces in a complex biological medium, in contrast to its shorter PSar analogues. These stealth NPs bearing HaloTag ligand enabled specific targeting of proteins at the cell surface. The obtained results show that a relatively short PSar peptide can be used for achieving stealth properties in polymeric NPs, allowing specific protein targeting with minimized non-specific interactions. The obtained PSar-functionalized polymeric NPs appear as a powerful platform for the fabrication of the next generation of nanomaterials for bioimaging and biosensing applications.

## Introduction

Nanoparticles (NPs) have gained significant interest in the last decades due to their unique properties and multitude of applications, especially in biology and medicine. It concerns, in particular, drug delivery and bioimaging applications, where the nanoparticle serves as a nano-vehicle that delivers biologically active molecules or contrast agents at the cellular or animal level.^1^ In this respect, luminescent nanoparticles attracted attention due to their exquisite brightness compared to organic dyes,^2^ multi-functionality in terms of load and targeting/recognition groups, and high surface-to-volume ratio compared to bulk materials.^3^ These unique properties make them particularly powerful for single-particle tracking, in vivo imaging in small animals, theranostics application to track the delivery of drugs, as well as biosensing applications for the detection of biologically important molecules with superior sensitivity.^4^ Luminescent NPs can be built from a variety of materials, and they could be split into inorganic NPs, such as semiconductor quantum dots,^5^ dye-doped silica NPs,^6^ metal clusters,^7^ metal-organic framework NPs,^8^ carbon dots,^9^ etc. and organic NPs,^10^ which include conjugated polymer NPs,^11-13^ aggregation-induced emission NPs,^14-16^ dye-loaded lipid^17^ and polymer^18-19^ NPs.

Despite an impressive variety of materials used for NP fabrication, the surface functionalization for biological applications is generally done by the same type of organic molecule – polyethylene glycol (PEG).^20-21^ Through the years, PEG has established a leading position as low-toxic material, with universal capacity to provide water solubility and biocompatibility to various nanomaterials. Moreover, PEG confers so-called stealth properties to NPs, which prevent interaction of the NPs surface with biomolecules, such as proteins of serum or cell membrane proteins.^22^ These stealth properties are of exceptional importance for targeted biological imaging or drug delivery applications because they minimize non-specific interactions and confer NPs long circulation time required for efficient targeting of diseased tissues.^23-25^ However, these decades of research accumulated evidences that PEG can cause a number of adverse biological effects, which raised significant concerns about its safety. ^26-30^ First, it was shown that the administration of PEGylated nanomaterials can cause an immunological response after repetitive administration. ^31-33^ Moreover, PEG is generally considered as a non-biodegradable polymer.^34^ Despite its well-developed chemistry, it is hard to obtain PEG of precise length. Finally, stealth properties of NPs are usually achieved with relatively long PEG chains of 3000-5000 Da. ^20-21,35^

To overcome these limitations, a number of PEG alternatives were developed.^30,36^ Thus, other synthetic polymers such as poly(glycerols) (PGs),^37^ poly(oxazolines) (POx), ^38-39^ poly(hydroxypropyl methacrylate) (PHPMA),^40^ poly(2-hydroxyethyl methacrylate) (PHEMA),^41^ poly(N-(2-hydroxypropyl)methacrylamide) (HPMA),^42^ poly(vinylpyrrolidone) (PVP),^43^ poly(N,N-dimethyl acrylamide) (PDMA),^44-45^ Poly(N-Methyl-N-Vinylacetamide),^46^ poly(N-isopropilacrylamide) (PNIPAM),^47^ and poly(N-acryloylmorpholine) (PAcM)^48^ and zwitterionic polymers, containing carboxybetaine,^49^ sulfobetaine,^50^ and phosphorylcholine^51^ groups, have also been used to replace PEG, and their synthesis and properties are reviewed elsewhere. ^29,52^ However, most of those molecules are built with highly stable C-C and C-N bonds in their backbone, making their biodegradability questionable.

Peptide-functionalized nanoparticles appear as promising platforms for advanced nanomaterials for biological and biomedical applications.^53-55^ The synthetic analogues of natural peptides are particularly promising alternatives of PEG, where the key example is polysarcosine (PSar), built of sarcosine (N-methyl glycine).^56^ The latter is a non-canonical naturally occurring amino acid, which is largely found in the muscles and other tissues. PSar is a hydrophilic and non-ionic polymer with properties similar to PEG.^57-59^ PSar offers additional advantages as it is biodegradable, non-toxic, and non-immunogenic. ^56,60^ Indeed, as it is built from endogenous (naturally present) amino acid, it is safe for human use. Moreover, PSar is truly a biodegradable polymer due to the presence of amide bonds, cleavable by enzymes. Molecular dynamics simulations showed that PSar exhibits stealth properties with respect to serum proteins.^61^ In literature, the polysarcosine chains of 50 to 200 repeating units have been employed to stabilize gold nanoparticles,^62-63^ quantum dots,^64^ and other inorganic NPs,^65^ as well as liposomes^66-67^ micelles,^68^ mRNA-lipid NPs,^69^ mRNA-polymer NPs,^70^ antibody-drug conjugates,^71^ etc. However, to the best of our knowledge, the effect of length for short PSar peptides on NPs stealth properties have not been studied to date. In one example, PSar with varied length from 23 to 68 was grafted to liposomes.^67^ These studies showed that the longest PSar ensured the longest circulation time and decreased immunogenicity compared to PEG. More work is needed to clarify whether shorter PSar units could also provide stealth properties to NPs.

In particular, it concerns polymeric NPs, which are powerful platforms for bioimaging and drug delivery.^72-73^ Peptide-functionalized polymeric NPs constitute an important research direction, which has yielded a variety of materials for biomedical applications. ^74-75^ However, the examples of polymeric NPs functionalized with PSar are still rare, ^70,76-78^ and the effect of PSar or the use of short PSar peptides is not documented. In the last decade, we focused our efforts on dye-loaded polymeric NPs, which emerged as powerful nanoscale tools for biosensing and bioimaging. ^2,19^ Their polymer matrix is an excellent reservoir for the encapsulation of dyes, other contrast agents, and drugs, while its surface remains versatile. These NPs can be obtained by nanoprecipitation of hydrophobic polymers bearing few charged groups that control particle size.^79-80^ Their loading with cationic dyes with bulky hydrophobic counterion ensured efficient encapsulation with minimal dye leaching and minimized aggregation-caused quenching^81-82^ yielding NPs of exceptional brightness.^83-84^ Their surface was successfully functionalized with oligonucleotides using strain-promoted cycloaddition (click) reaction, which yielded ultrasensitive probes for DNA and RNA detection,^85-86^ important for diagnostics applications.^87-88^ The use of zwitterionic groups could improve their stealth properties.^50^ These NPs functionalized with biotin enabled grafting streptavidin and biotinylated antibody for specific targeting.^89^ However, so far, these types of NPs have not been functionalized with peptides. The use of PSar for their functionalization could yield new nanomaterials with enhanced stealth and targeting properties.

In the present work, functionalized dye-loaded polymeric NPs with PSar of different lengths ranging from 5 to 19 sarcosine units. The obtained peptide-functionalized NPs showed remarkable colloidal stability in physiological media. The length of PSar showed a profound effect of NPs properties: the higher PSar lengths led to a nearly neutral surface charge and minimized interactions with serum proteins, as evidenced by fluorescence correlations spectroscopy. The NPs with the longest PSar (19 units, molecular weight of ∼1400 Da) showed minimal interactions with the cells and glass surfaces in a complex biological medium compared to its shorter PSar analogues. These were further applied for specific targeting of proteins at the cell surface using HaloTag strategy. The obtained results show that a relatively short PSar peptide can be used for achieving stealth properties in polymeric NPs, allowing specific protein targeting with minimized non-specific interactions.

## Results and discussion

### Design and synthesis

Dye-loaded polymeric NPs were based on PEMA-MA polymer derivative. The PEMA-MA polymer was functionalized with aspartic acid-azide (Asp-N3), which presents charged carboxylate and reactive azide groups in close proximity. The charged carboxylate facilitates the formation of small NPs and ensures azide groups are exposed on the NP surface (NPs-N3), which was previously used for grafting oligonucleotides bearing dibenzocyclooctyne (DBCO) by stain-promoted cycloaddition reaction.^86^ Here, this approach was used for bio-conjugation of DBCO-polysarcosine glycine (DBCO-GSar) peptides on the surface of NPs, resulting in peptide-functionalized NPs (NPs-GSar) (Figure 1). The peptide shell is expected to provide a stealth layer around the NPs, enhancing their stability in a native cellular environment and minimising non-specific interactions, thereby enabling bioimaging applications. Here, polysarcosine glycine chains with 6, 12, and 20 repeating units were synthesized to achieve stealth properties while minimizing chain length. The peptides were synthesized using an automated solid phase peptide synthesizer with a Glycine-Wang resin. Following cleavage from the resin, the peptides were conjugated at their N-termini with DBCO-NHS ester (methods and materials, section 2.3). This resulted in the formation of DBCO-GSar peptides of three different chain lengths: GSar5, GSar11, and GSar19 for further grafting to NPs-N3. The purity and identity of the conjugates were confirmed by HPLC and mass spectrometry, respectively (methods and materials, section 2.5 & 2.6).

**Figure 1.**
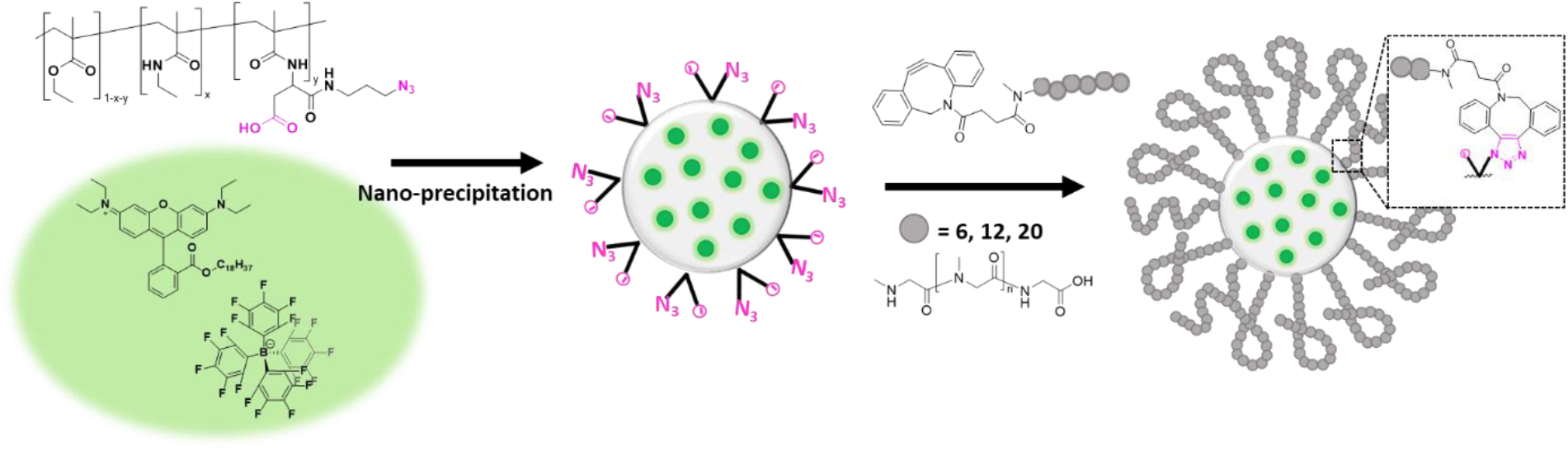
Scheme for synthesis of peptide-functionalized dye-loaded fluorescent polymeric nanoparticles. Nanoprecipitation of polymer PEMA-AspN3 with fluorescent dye to obtain NPs bearing azide groups, peptide (DBCO-GSarn) functionalization on the surface of NPs-N3. Chemical structures of PEMA-AspN3 polymer, rhodamine derivative R18, its bulky counter ion F5-TPB and DBCO-GSarn peptides are also shown.

### PSar-functionalized polymeric nanoparticles

To prepare NPs-N3, we first synthesized a conjugate of PEMA polymer with Asp-N3, but we changed the approach compared to our previous report.^86^ We first synthesized PEMA polymer bearing an activated carboxylic acid with a pentafluorophenyl group (5%), which was subsequently coupled with a previously synthesized linker Asp(OtBu)-N3.^86^ NMR spectra revealed a good yield of polymer modification (56%). Finally, the tert-butyl group was removed from the polymer to afford the final product PEMA-AspN3. Nanoprecipitation of polymer PEMA-AspN3, along with fluorescent dye ion pair (R18/F5-TPB) into an aqueous buffer (pH 7.4) gave NPs of 28 nm diameter according to transmission electron microscopy (TEM). The NPs were loaded with 9 wt% of R18/F5-TPB dye (vs total mass of NPs), and the fluorescence quantum yield of the NPs measured was 0.15 ± 0.015 (n = 3) (Figure S1).

For peptide grafting on the NP surface, the obtained dye-loaded NPs-N3 were reacted with DBCO-GSarn peptides (GSar5, GSar11, and GSar19) at 40°C for 22 h, giving three peptide-functionalized NPs: NPs-GSar5, NPs-GSar11, and NPs-GSar19, respectively. Purification of these peptide-functionalized NPs from non-reacted peptides was achieved by successive ultrafiltration through 100kDa Amicon filters. The successful peptide functionalization was confirmed by the increase in NP size and stability against the filters during ultrafiltration. According to TEM, the obtained three peptide-coated NPs were spherical, monodisperse, and showed no signs of aggregation. The diameter measured by TEM were 34.6±4.7 nm for NPs-GSar5, 35.9±3.5 nm for NPs-GSar11, and 34.1±4.9 nm for NPs-GSar19, compared to 28.6±4.1 nm for NPs-N3 (Figure S2a, S2b, 2b, 2a). Dynamic light scattering (DLS) measurements showed a systematic increase in the hydrodynamic diameter of peptide-coated NPs and good PDI (Table 1). The size increase was correlated with the increase in the peptide chain length (GSar19> GSar11> GSar5). Additionally, the zeta potential of NPs (Table 1) shifted from strongly negative for NPs-N3 (−33 mV) to slightly negative for NPs-GSar5 and NPs-GSar11 (−10.5 and -6.6 mV, respectively) and nearly neutral for NP-GSar19 (−2.1 mV). This result indicates an effective shielding of the negative charge of the carboxylate ions by the relatively neutral polysarcosine glycine peptides (Table S1). Similar values have been reported by Chen *et. al*. for gold NPs coated with polysarcosine brushes.^62^ The charge neutralization after peptide coating is expected to prevent protein corona, as a neutral charge should minimize interactions with proteins.

**Table 1.** Hydrodynamic diameter (nm) and zeta potential (mV) for nanoparticles NPs without modification and modification with three different chain lengths of peptides.

| Sample | Hydrodynamic radius (nm) | Polydispersity index (PDI) | Zeta potential (mV) |
| --- | --- | --- | --- |
| NPs-N3 | $34.5 \pm 1.9$ | 0.09 | $-33.2 \pm 5.1$ |
| NPs-GSar5 | $36 \pm 0.8$ | 0.11 | $-10.5 \pm 1.3$ |
| NPs-GSar11 | $38 \pm 0.5$ | 0.08 | $-6.63 \pm 0.55$ |
| NPs-GSar19 | $39 \pm 1.0$ | 0.07 | $-2.18 \pm 0.55$ |

<sup>a</sup> Error is s.d.m. from three repeated experiments; <sup>b</sup> error is s.d.m. of three sequential measurements of the same sample.

Peptide-functionalization on NPs surface was confirmed by stability towards ultrafiltration through 100 kDa filter, where the hydrodynamic diameter of NPs-GSar remained consistent after each round of ultrafiltration (Figure 2d, S2c, S2d). In contrast, bare NPs-N3, showed a drastic increase in the diameter and a high polydispersity index, indicating aggregation of NPs (Figure 2c). UV-visible absorption spectra were also recorded after each round of ultrafiltration. For bare NPs-N3, a decrease in the absorbance at the maximum and the recovery efficiency of only 44% (Figure 2e) were observed, which suggests that they aggregated and stuck on the filter during ultrafiltration. However, after peptide-functionalization, the efficiency of NPs recovery improved to approximately 70% (Figure 2f, S2e, S2f). Overall, the increase in hydrodynamic diameter of the peptide-functionalized NPs, change in their zeta potential, and their higher colloidal stability towards the ultra-centrifugal filters validates the successful conjugation of peptides on NPs surface.

**Figure 2.**
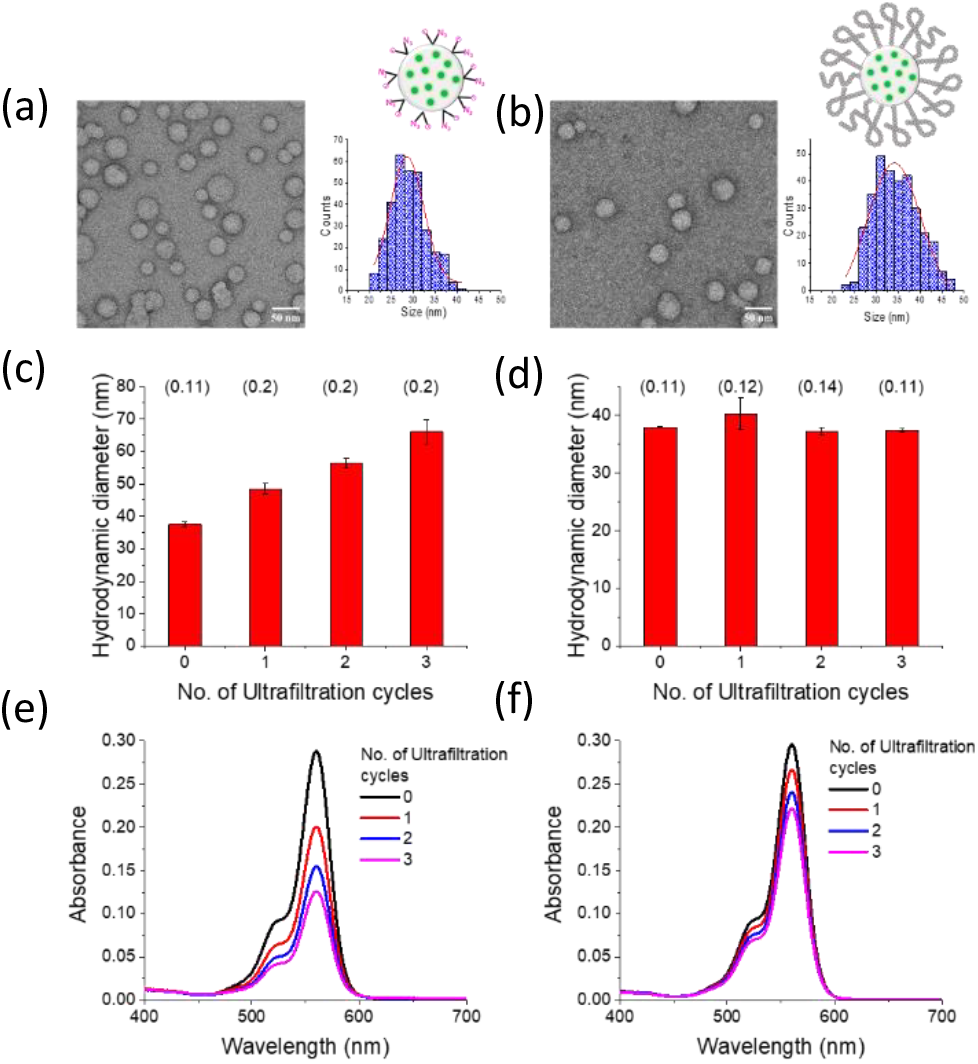
Characterisation of peptide-functionalized NPs. TEM images of (a) NPs-N3 and (b) NPs-GSar19 and their corresponding size distribution statistics. Scale bar 50 nm. Size by DLS of NPs after each round of filtration and their polydispersity index (PDI) mentioned in brackets for (c) NPs-N3 and (g) NPs-GSar19. The absorbance spectra for (e) NPs-N3 and (f) NPs-GSar19 after each round of ultrafiltration.

### Stealth properties of peptide-functionalized NPs

The colloidal stability of peptide-functionalized NPs was evaluated in the presence of salt (NaCl) at a physiological concentration (150 mM), where the size of the NPs was measured by Fluorescence Correlation Spectroscopy (FCS). The presence of salt caused an increase in NPs size of NPs-N3, probably due to their aggregations (Figure 3). In contrast, the peptide-functionalized NPs showed no signs of aggregation or precipitation, indicating that the polySar shell conferred stability. Additionally, no significant changes in the fluorescence brightness were detected for NPs-GSar in the presence of salt, confirming the colloidal stability of NPs in PBS (Table S1, S2). The influence of peptide chain lengths stealth properties of NPs was studied by FCS in the presence of 5% fetal bovine serum (FBS), a complex mixture containing notably serum albumin and other blood proteins. In the case of uncoated NPs-N3 and NPs-GSar5, an increase in size of 15-20 nm was observed, probably due to the formation of protein corona and aggregation of NPs. NPs-GSar11 showed a smaller increase in size (∼4 nm), indicating some adsorption of serum proteins on the NPs surface but without NP aggregation. Remarkably, no increase in size was observed for NPs-GSar19, suggesting the longer peptide chains effectively protected NP surface from protein adsorption (Figure 3, S3). Thus, we found that longer polysarcosine (19-mer) chains conferred NPs-GSar19 stealth properties against serum proteins, while maintaining colloidal stability. To the best of our knowledge, there were no reports showing that stealth properties of polymeric NPs can be achieved with such a short polySar peptide, corresponding to the molecular weight of ∼1400 Da.

**Figure 3.**
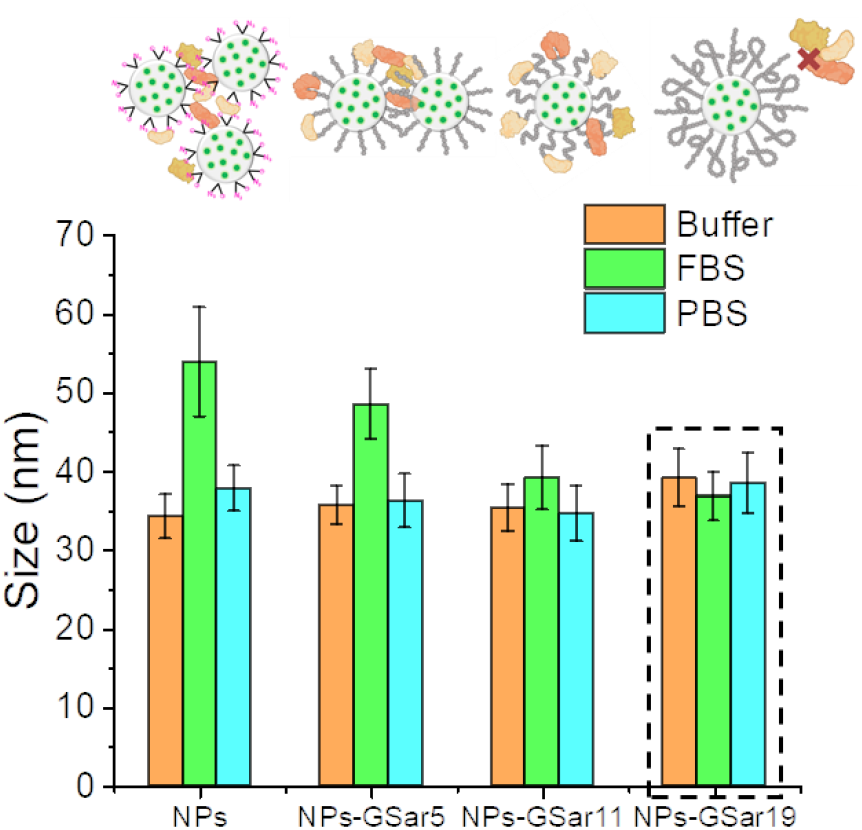
Size of NPs (non-modified and modified with 3 different peptides) by Fluorescence Correlation Spectroscopy (FCS) in buffer (orange), in the presence of 5% fetal bovine serum (green), and in phosphate buffer saline (Blue).

### Interaction of peptide-functionalized NPs with glass surface

We investigated the non-specific interaction of NPs with a glass surface using fluorescence microscopy in three different commercially available physiological media. We were interested in finding optimal conditions where NPs-GSar19 does not adhere to the glass surface, thus minimizing background noise. Significant non-specific adsorption of NPs to the glass surface was observed in the presence of HBSS and PBS media, even though peptide-functionalized NPs showed a tendency to adsorb less. However, in Opti-MEM media, the adsorption of all peptide-functionalized NPs was dramatically reduced. We suppose that this nutrient-rich medium containing vitamins, growth factors, some proteins, etc may create a protective organic layer on the glass surface, which substantially decreases the adsorption of peptide-functionalized NPs to the glass surface. In particular, NPs-GSar19 stood out showing no signal of NPs on the glass surface in Opti-MEM, indicating no interaction with the glass surface (Figure 4a, S4a). These findings suggest that NPs-GSar19 not only show stealth properties in serum, but also show minimized non-specific interactions with the surface. However, the media composition is crucial in reducing non-specific interaction and background noise even for these stealth NPs.

**Figure 4.**
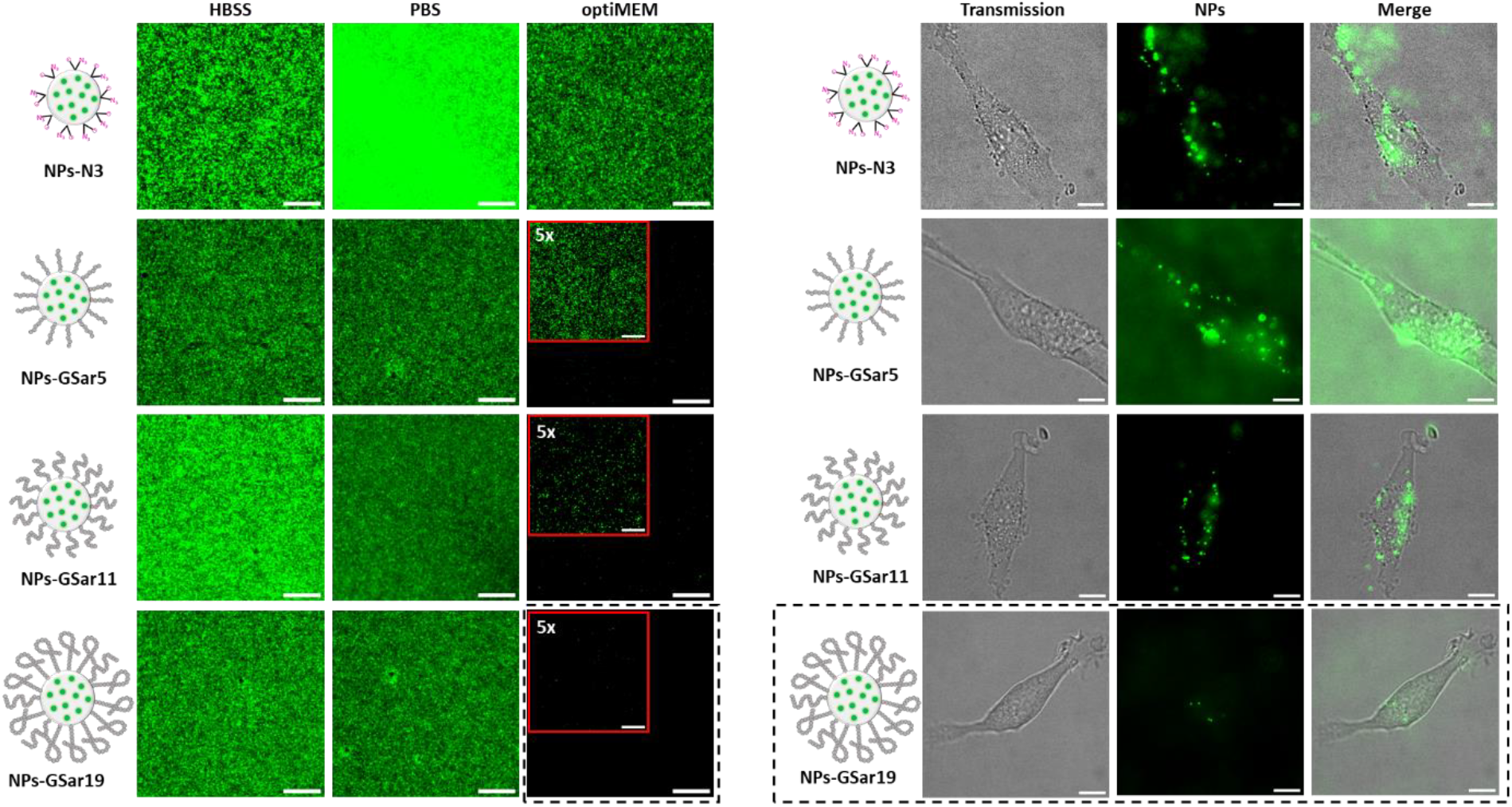
(a) Non-specific labelling of NPs-N3 (20 pM) and NPs-GSarn (20 pM) with glass surface using epi-fluorescence microscopy in 3 different biological medias (i) Hank’s balanced salt solution (HBSS), (ii) Phosphate buffer saline (PBS) and (iii) optiMEM. The B/C is 1000-6000. Scale bar 10 μm. (b) Non-specific labelling of cell surface of live U87 cells using epi-fluorescence microscopy with NPs-N3 (20pM) and NPs-GSarn (20 pM). B/C is 20-500. Scale bar 10 μm. The black dotted line shows the best conditions for stealth’s NPs.

### Interaction of peptide-functionalized NPs with cell surface

The surface modification of NPs with peptides may alter their non-specific interactions with the cell surface, which is essential for their specific targeting. The effect of peptide-functionalization on the NPs-cell surface interaction was evaluated by fluorescence microscopy using U87 cells in optiMEM media, which showed reduced background noise. The U87 cells were incubated with unmodified and peptide-modified NPs for 20 min, and the extent of non-specific labelling was studied by an epi-fluorescence microscope. A trend of decreasing cell surface labelling by NPs with increased peptide chain length was observed. Thus, bare NPs-N3 and NPs-GSar5 exhibited the strongest fluorescence signal, indicating the highest non-specific interactions with cells, whereas NPs-GSar19 showed the lowest signal (Figure 4b, S4b). This reduced interaction was attributed to stealth properties of GSar19 peptide, which effectively shielded NPs surface, prohibiting non-specific interactions with proteins of the cell surface.

The interactions of NPs with cells were then quantified using Flow cytometry. The signal from NPs-GSar19 was lower than that of NPs-GSar5 (Figure S7). One should note that when the labelling experiments were conducted in the presence of 5% FBS, overall non-specific labelling of NPs was reduced, and the difference within different NPs was minimized. However, the trend of cell surface labelling with NPs remains consistent, with NPs-GSar19 showing the least interaction with the cell surface (Figure S5-S7). Thus, NPs functionalised with GSar19 peptide was clearly the best candidate for stealth NPs, exhibiting stability in the presence of salts, and serum proteins, negligible adsorption on glass surface, and minimized non-specific interaction with U87 cells. These properties highlighted the potential of NP-GSar19 as a suitable candidate for specific targeting applications with minimized background noise.

### Peptide-functionalized NPs for HaloTag targeting

To further demonstrate the utility of our stealth NPs in targeting and bioimaging applications. We focused on targeting proteins bearing genetically encoded protein tags, specifically HaloTag,^90^ which is a universal tool for specific protein labelling. The haloalkane dehalogenase-based HaloTag, is a widely used protein tag that enables covalent ligation to chloroalkane ligands, providing specific and robust labelling under physiological conditions. Thus, we designed the peptide-functionalized NPs presenting chloroalkane ligand at the peptide extremity (NPs-GSar19-Cl, Figure 5a). For that, the DBCO-GSar19 peptide was modified at its carboxyl end with a chloroalkane ligand to yield DBCO-GSar19-Cl peptide. The obtained DBCO-GSar19-Cl and the non-modified stealth peptide (DBCO-GSar19) at 1 mol% or 10 mol% of the former were grafted together to NPs-N3 (Figure 5a). According to TEM, NPs-GSar19-Cl showed monodispersed, spherical NPs with a size of 32±4 nm (Figure S2). The successful peptide functionalization was confirmed by the stability of NPs towards ultra-centrifugal filters, where the hydrodynamic diameter of NPs remained consistent, and the absorbance spectra indicated ∼50 % of NPs were recovered. Further analysis by FCS confirmed the stealth behaviour of NPs-GSar19-Cl, with the NPs size remaining stable in the presence of 5% FBS and PBS (Figure S8). Next, we studied the specific ligation of NPs-GSar19-Cl with HaloTag-green fluorescent protein (HT-GFP), as compared to green fluorescent protein (GFP) as a reference (Figure 5b). Both proteins were incubated with NPs-GSar19 containing 1 mol% of targeting ligand DBCO-GSar19-Cl, and then the non-reacted protein was removed via ÄKTA size-exclusion chromatography. Absorption and fluorescence spectra were recorded before and after purification for each separated fraction to access ligation specificity. Fluorescence spectra recorded at an excitation wavelength of 470 nm showed two emission peaks; one at 510 nm corresponding to GFP and another at 580 nm for NPs. After purification, three chromatographic fractions (T11, T12, and T13) exhibited absorbance maximum at 560 nm, indicating the presence of NPs (Figure S9 and S10). Among them, aliquot T12 showed the highest concentration of NPs (Figure S10) and was selected for further analysis. After purification, no emission maxima at 510 nm was observed for GFP without HaloTag, suggesting that it was completely removed after purification of NPs. In contrast, HT-GFP retained its emission maximum at 510 nm, indicating its successful immobilization on the NPs-GSar19-Cl and confirming the specificity of the system to HaloTag (Figure 5c). Furthermore, the size of purified NPs determined by FCS increased by ∼12 nm when reacted to HT-GFP, whereas no significant size change was observed for NPs incubated with GFP. A small increase of ∼4 nm in size for the latter was attributed to peptide functionalization (DBCO-GSar19/ DBCO-GSar19-Cl) onto the NPs (Figure 5d). Our result suggested the selective ligation of HaloTag proteins to NPs-GSar19-Cl.

**Figure 5.**
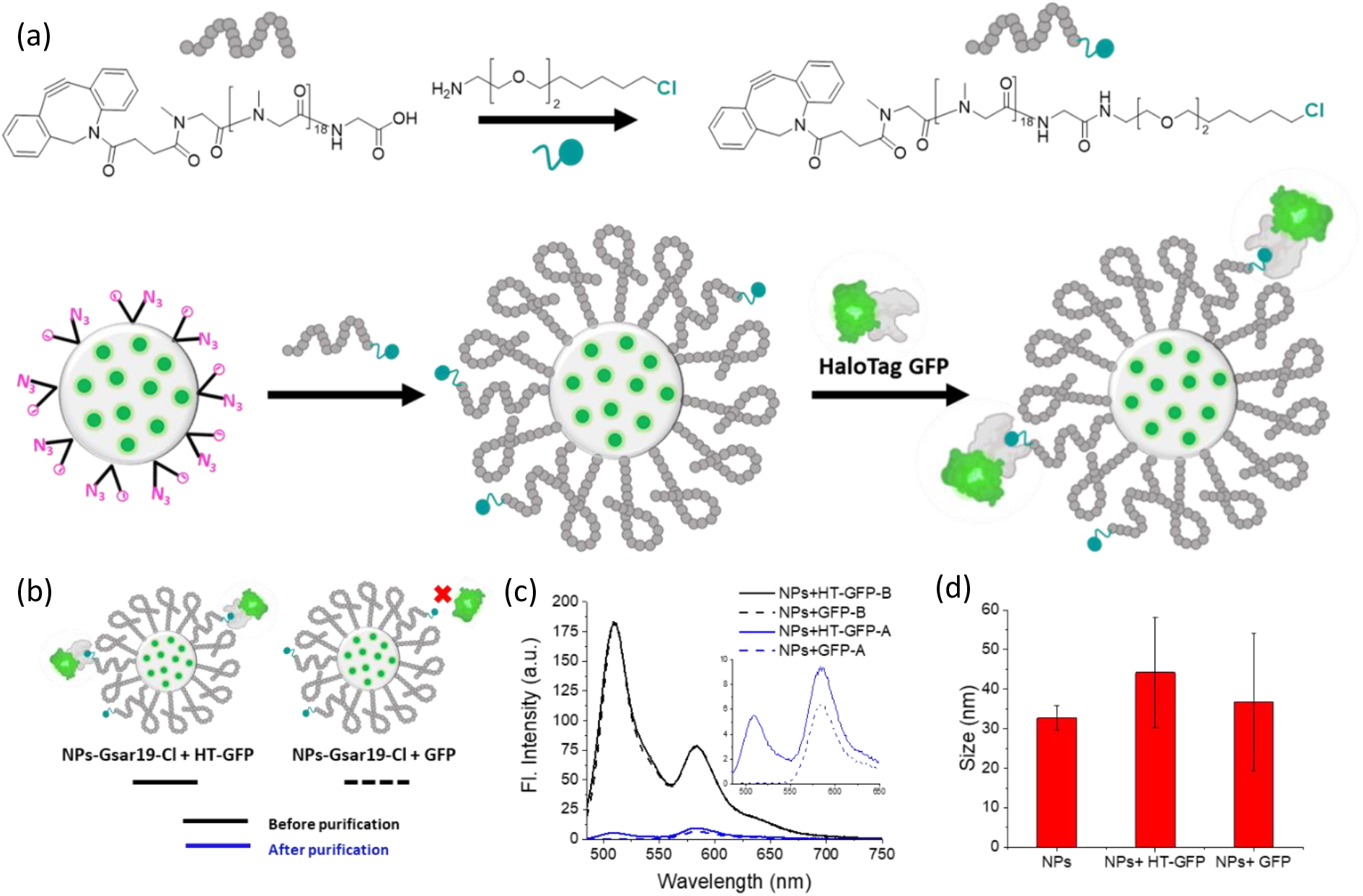
(a) Scheme for synthesis of chloroalkane ligand modified Stealth NPs (NPs-GSar19-Cl) for labelling HaloTag GFP protein. Structure of Stealth peptide DBCO-Gsar19 modified with chloride ligand at C-terminal, which was immobilized on NPs-N3. (b) illustration showing the specific immobilisation of HaloTag GFP as compared to GFP on NPs-GSar19-Cl. (c) The fluorescence intensity of HaloTag-GFP protein (Solid line) and GFP protein (dash line) immobilized on NPs-GSar19-Cl before (black) and after (blue) AKTA purification excited at 470 nm. (d) Size of NPs by FCS for NPs-N3; and NPs-GSar19-Cl with immobilised HaloTag GFP and NPs-GSar19-Cl with GFP.

We quantified the number of HT-GFP immobilized per NP by plotting a calibration of observed fluorescence intensity maxima versus the concentration of HT-GFP. It was found that 10 HT-GFP and 36 HT-GFP were immobilized per NP-GSar19-Cl having 1% and 10% of chloroalkane ligands in the T12 fraction, respectively (Figure S10 & S11). Herein, a higher number of chloroalkane ligands were used to maximise the potential for HaloTag protein labelling in cellular applications.

### HaloTag protein targeting in cells

Finally, we explored the labelling of HaloTag proteins in complex biological media with NPs-GSar19-Cl bearing 10% of chloroalkane ligand. HEK293T cells were transfected with plasmid DNA pAG842,^91^ encoding for HaloTag protein fused to platelet-derived growth factor receptor (PDGFR), expressed on the outer cell surface (see Materials and Methods, section 1.6). A 200 pM concentration of NPs-GSar19-Cl was employed to label the cell surface, while non-specific interactions were minimised with the help of serum proteins (5% FBS) (Figure S12) or surfactants (0.01 g/L of Tween20) (Figure S13). We observed excellent labelling of cell surface for transfected cells expressing HaloTag protein (Figure 6A). The cell surface labelling with NPs co-localized well with the commercially available Janelia Flour-650 bearing chloroalkane ligand (Figure 6A). In contrast, no labelling was observed on the cell surface of non-transfected cells. Quantitative image analysis confirmed large differences between the signals recorded from transfected and non-transfected cells, thus validating the selectivity of NPs-GSar19-Cl for HaloTag protein-expressing cells. This result highlights the potential of stealth peptide-functionalized NPs for specific targeting in cellular applications.

**Figure 6.**
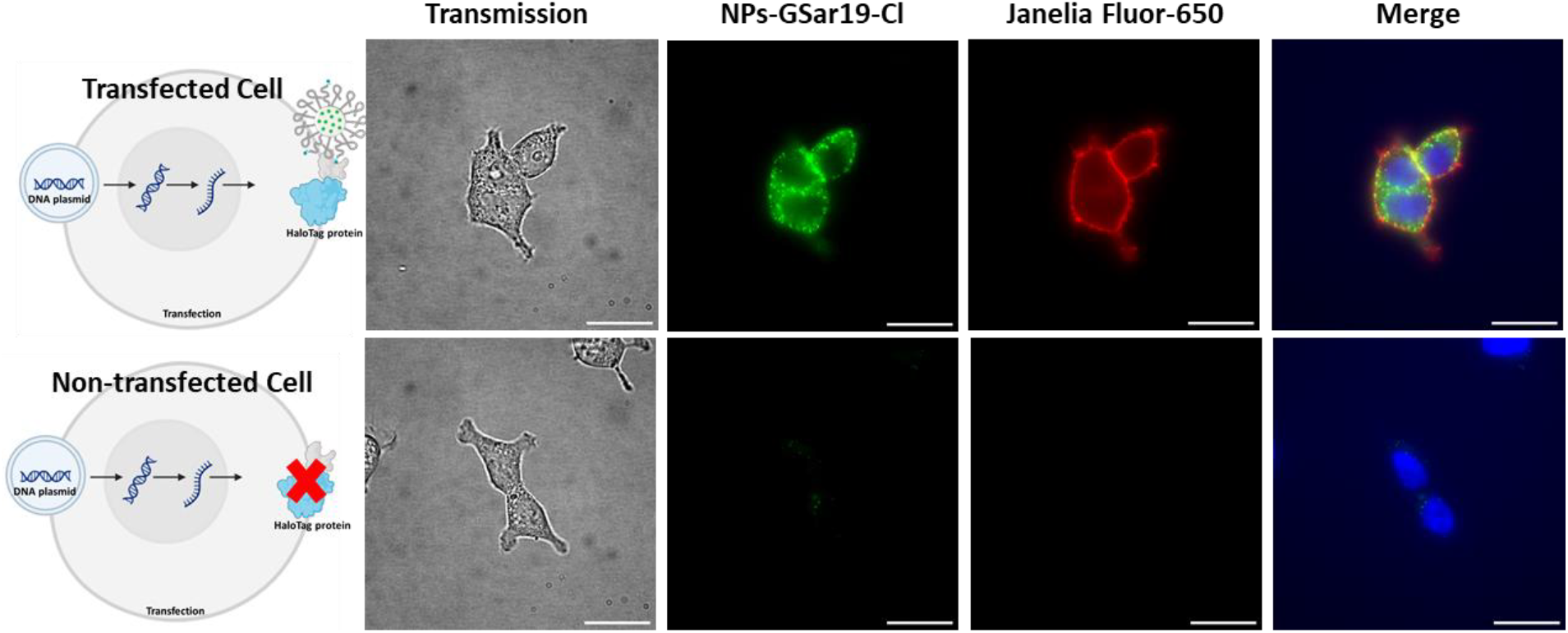
Microscopy images for labelling of HaloTag proteins with 200 pM NPs-GSar19-Cl in presence of 5% FBS (green colour) and 1 nM commercial Janelia Fluor-650 dye (red colour) expressed on cell surface of transfected HEK293T cells (upper panel) and non-transfected HEK cells (lower panel). The nucleus was labelled with Hoechst dye in blue colour. B/C for green channel 30-650 and 100-5000 for red channel. Scale bar 25 μm.

## Conclusions

Stealth properties of nanoparticles prevent them from non-specific interactions with biomolecules and cells, which is crucial for their applications for specific targeting and imaging and in biological systems. The commonly used PEG reaches its limits as stealth shell of NPs, because of some immunogenicity, non-biodegradability, relatively high length required for achieving stealth properties. As a promising replacement of PEG, we considered polysarcosine (PSar), which has already shown its stealth properties in applications to some nanoparticles. Nevertheless, relatively long PSar polypeptides were used for this purpose, so it remains unclear about the role of PSar length and the minimal size required to achieve stealth properties of NPs. Moreover, application of PSar to polymeric NPs remains rare and requires additional research given raising importance of polymeric nanomaterials in biological and biomedical applications. In this work we introduce peptide-functionalized dye-loaded polymeric NPs, which explore PSar of different lengths with the aim to obtain stealth NPs. Polymeric PEMA-based NPs loaded with rhodamine R18 dye with bulky hydrophobic counterion F5-TPB and bearing azide groups at their surface were functionalized with PSar of different lengths ranging from 5 to 19 sarcosine units using strain-promoted cycloaddition reaction. The peptide grafting resulted in a small increase in the particle hydrodynamic diameter, the signifying drop in the negative zeta-potential and increase in the particle stability in the conditions of ultra-filtration. The obtained peptide-functionalized NPs showed remarkable colloidal stability in physiological media. The length of PSar showed a profound effect on stealth properties of NPs. Indeed, the increase in the length of grafted PSar lead to decreased protein adsorption according to fluorescence correlations spectroscopy. The NPs with 19mer PSar showed minimal interactions with live cells and glass surfaces in a complex biological medium, in contrast to its shorter PSar analogues, such as 5mer and 11mer. These stealth NPs bearing HaloTag ligand enabled specific targeting of proteins at the cell surface, where control cells without protein of interest showed negligible signal. It is important to stress that the PSar19 used has a molecular weight of 1400 Da, which is lower than that required for PEG polymers to achieve stealth properties (2000 to 5000 Da). It is also much lower than that used in the previous studies of PSar-functionalized NPs (50-100 sarcosine units). This means that a relatively short PSar peptide allows achieving stealth properties in polymeric NPs and enable operation in complex media for specific protein targeting on the cell surface with minimized non-specific interactions. This work introduces peptide-functionalized polymeric NPs based on PSar as a perspective nanoscale platform for the fabrication functional nanomaterials for bioimaging and biosensing applications.

## Supporting information

Supplemental information for publication

## Acknowledgements

This work was supported by Agence Nationale de la Recherche (ANR) AmpliSens ANR-21-CE42-0019-01. We acknowledge the PSSP platform of University of Strasbourg for the peptide synthesis work. For Electron microscopy, this work used the Integrated Structural Biology platform of the Strasbourg Instruct-ERIC center IGBMC-CBI supported by FRISBI (ANR-10-INBS-0005).

