## Supplemental information for publication for "Peptide-functionalized fluorescent polymeric nanoparticles: polysarcosine length defines stealth properties"

**Table of contents**

**Methods and materials**

1. General materials and methods
   - 1.1 Materials
   - 1.2 HPLC
   - 1.3 Nanoparticle Characterization
   - 1.4 Transmission electron microscopy
   - 1.5 Fluorescence correlation spectroscopy
   - 1.6 Cell lines, culture and treatment
   - 1.7 Fluorescence microscopy
   - 1.8 Flow cytometry
   - 1.9 ÄKTA purification
2. Synthesis and characterization of polymer & peptides
   - 2.1 Synthesis of polymer
   - 2.2 Synthesis of peptides
   - 2.3 Modification of peptides with DBCO (DBCO-acid to DBCO-NHS ester)
   - 2.4 Modification of peptide with HaloTAG.
   - 2.5 HPLC traces
   - 2.6 NMR and Mass Data
   - 2.7 Preparation of dye-loaded nanoparticles
   - 2.8 Synthesis of peptide-functionalized Nanoparticles
   - 2.9 Synthesis of chloroalkane ligand modified peptide-functionalized nanoparticles
   - 2.10 Immobilization of HaloTag GFP on NPs-GSar5-Cl
   - 2.11 Molecular Biology
   - 2.12 Protein Expression and Purification

**Supporting figures**

**Supporting Tables**

**References**

### 1. General materials and methods

#### 1.1 Materials.

Chemical compounds: Fmoc-Sar-OH (HPLC, >98%) was purchased from BLD Pharma. Oxyma (HPLC, .99%), and *N,N'*-Diisopropylcarbodiimide (98%, for peptide synthesis) were purchased from TCI. Piperidine (for synthesis), Triisopropylsilane (for synthesis), and Dibenzocyclooctyne-carboxylic acid were purchased from Sigma-Aldrich. Acetonitrile (HPLC, 99.8%), dichloromethane (for analysis, ≥99.8%), Diethylether (99%), Dimethylformamide (99.5%) were purchased from FischerScientific. Amicon Centrifugal filters (0.5mL, 100KDa) was purchased from Merck milipore. Fmoc-Gly-Wang resin (0.34mmol/g loading) was purchased from Activotec. Trifluoroacetic acid (99%) was purchased from Alfa Aesar. Sodium phosphate monobasic (>99.0%, Sigma-Aldrich) and sodium phosphate dibasic dihydrate (>99.0%, Sigma- Aldrich) were used to prepare 20 mM phosphate buffers at pH 7.4. Milli-Q water (Millipore) was used in all experiments. µ-dish ibidi were purchased from ibidi and LabTek chamber (borosilicate glass, eight-wells) was purchased from ThermoFischer Scientific for microscopy.

#### 1.2 HPLC: Agilent Infinity 1260 II

The detection and purification of peptides and their modification using HPLC was performed on Agilent infinity 1260 II system with UV detector. 210 nm and 310 nm wavelengths were used for detection of DBCO-modified peptides. The method used reverse-phase chromatography with Agilent Poroshell 120 C18 4 µm column for analysis and Uptisphere Strategy C18 5 µm 250 × 10 mm semi-preparative LC column for purification of peptides. A gradient flow of solution A (water + 0.01% TFA) and Solution B (Acetonitrile + 0.01% TFA) was used. For analytical run, 0-3 min 100% A; 3-17 min gradient of solution A to 0%, 17-18.5 min 0% A and 18.5-20 min 100% A. The mobile phase flow rate was 1 ml/ min. And for Semi-preparative column, 0-5min 100% solution A; 5-30 min gradient of solution A to 0%, 30-35 min 0% A and 35-40 min 100% A gradient. The mobile phase flow rate was 2 ml/ min.

#### 1.3 Nanoparticle characterisation

Measurements for the nanoparticle size and zeta potential were performed on a Zetasizer Nano ZSP (Malvern instruments S.A.). The mean value of the hydrodynamic diameter of the size distribution per volume was used for analysis. For zeta potential, three successive measurements combining electrophoretic mobility and laser Doppler velocimetry with >10 runs each were performed with an applied voltage of $\pm 150\,V$. Absorption spectra were recorded on a Cary 4000 UV-visible spectrophotometer (Varian), emission and excitation spectra were recorded on a FluoroMax-4 spectrofluorometer (Horiba Jobin Yvon). For standard fluorescence spectra the excitation was set to

530 nm. The fluorescence spectra were corrected for detector response and lamp fluctuations. The steady state spectra were recorded in buffer for NPs.

Quantum yields[1] of donor dye in NPs were calculated using R101 in methanol as a reference dye (Q.Y. = 1.0) with an absorbance of <0.1 at 530 nm.

#### 1.4 Transmission Electron microscopy

Carbon-coated coper-rhodium electron microscopy grids with 300 mesh (Euromedex, France) were surface treated with glow discharge in amylamine atmosphere (0.5 mbar, 4-5 mA, 30 s) in an Elmo glow discharge system (Cordouan Technologies, France). Then, 5 µL of NPs at 0.04 g $L^{-1}$ were deposited on the grids and left for 2 min in presence of 150mM NaCl. This step was repeated twice. The grids were then treated for 1 min with 2% uranyl acetate solution for staining. After that, the experiments were performed on a Philips CM 120 transmission electron microscope equipped with LaB6 filament and operating at 100 kV. Images were analysed using Fiji software. At least 150 particles per condition were analysed.

#### 1.5 Fluorescence correlation spectroscopy

Measurements were performed on a home-built confocal setup based on a Nikon inverted microscope with a Nikon 60× 1.2 NA water immersion objective. Excitation was provided by a continuous wave 532 nm laser diode (Oxxius) and photons were detected with a fibered avalanche photodiode (APD SPCM-AQR-14-FC, PerkinElmer) connected to an online hardware correlator (ALV7000-USB, ALV GmbH, Germany). Tetramethylrhodamine (Merck) was used as a reference for 532nm laser. The laser power used was 0.8 mW. The data was analysed using the PyCorrFit software in Triplet + 3D model fitting.

#### 1.6 Cell lines, culture and treatment

U87 (ATCC CCL-17) cells were grown in Eagle's Minimum Essential Medium (EMEM, Gibco Invitrogen) and supplemented with 10% fetal bovine serum (FBS, Lonza), 1% L-glutamine (Sigma-Aldrich), 1% nonessential amino acid solution (Gibco Invitrogen), and sodium pyruvate 1 mmol/L and HEK293T (ATCC CRL-3216) cells were grown in Dulbecco's modified eagle medium (DMEM, Gibco Invitrogen) and supplemented with 10% fetal bovine serum (FBS, Lonza), 1% L-glutamine (Sigma-Aldrich), 1% non-essential amino acid solution (Gibco, Invitrogen), and sodium pyruvate 1 mmol/L at 37 °C in a humidified 5% $CO_2$ atmosphere. Cells were seeded on a covered chambered glass coverslip (µ-dish, IBIDI) at a density of $8 \times 10^4$ cells/ well.

Transfection of HEK cells: HEK293T cells were seeded in ibidi chamber at a cell density of $1\times 10^5$ cells/ ibidi 24 h before transfection. Transfection mixture was prepared containing jetOPTIMUS buffer 200 µL and pDNA (0.5 µg/ mL) 2 µl was added. The solution was vortex and spin down. To it, jetOPTIMUS reagent was added 2 µL in a ratio of 1:1. The mixture was incubated for 10 min at r.t. before being added to cells. Cells were incubated in 1 mL DMEM media containing transfection mixture at 37 ℃ for 4-5 h. Then, the media was removed and cells were washed thrice with growing media DMEM and incubated again at 37 ℃, 5% $CO_2$ atm for 24 h before imaging.

Plasmid pAG842, encoding HaloTag fused to platelet derived growth factor receptor (PDGFR) transmembrane domain was used and was synthesized previously.[2]

### 1.7 Fluorescence microscopy

Single particle measurements were performed in the epi-fluorescence mode using Nikon Ti-E inverted microscope with 60x objective (oil, NA, Nikon). The excitation was provided by light emitting diodes (SpectraX, Lumencor) at 550 nm. The power density determined by power meter for 550 nm was 0.465 mW $cm^{-2}$. The fluorescence signal was recorded with Hamamatsu Orca Flash 4 camera. The exposure time was set to 300 ms per image frame.

### 1.8 Flow cytometry

Flow cytometry was performed on MACSQuant VYB (Miltenyi Biotec). Gates were defined to detect live cells by forward scattering (FCS) and side scattering channels (SSC). At least 150,000 events were collected for each condition for analysis using FCSalyzer software. 561 nm laser was used to excite R18 dye-loaded NPs and emission was recorded through 651/671 nm band pass filter.

### 1.9 ÄKTA purification

All purification of green fluorescent protein (GFP) immobilisation on NPs-GSar19-Cl were performed on ÄKTA start system (Cytiva). The ÄKTA system was equipped with rotatory type manual valve with sample injection through loop. The system column included a size exclusion column Sephacryl S200 200/10. The samples were analysed by single UV wavelength at 280 nm. The entire system was controlled by the software supplied by the manufacturer *UNICORN start* (Cytiva).

For the purification 20 mM Phosphate Buffer (pH 7.4) was used with a flow rate of 0.8 mL/min. The pressure limit was set to 0.8 bar. 0.5 mL of sample was injected in the loop.

### 2. Chemical Synthesis

NMR spectra were recorded at 20 °C on Bruker Avance III 400MHz spectrometer and chemical shifts were reported as delta scale in ppm relative to $CHCl_3$ (δ = 7.26 ppm) for $^1H$ NMR and $CDCl_3$ (δ = 77.16 ppm) for $^{13}C$ NMR. Mass spectra were obtained using an Agilent Q-TOF 6520 mass spectrometer.

#### 2.1 Peptide Synthesis

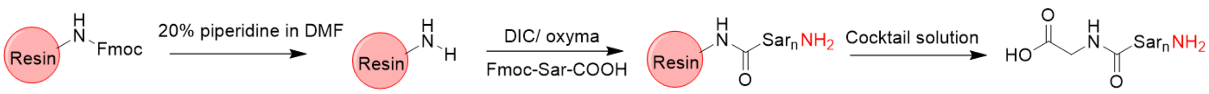


The polysarcosine glycine peptides were synthesized on a 0.1 mmol scale using fluorenylmethoxycarbonyl (Fmoc) chemistry on automated initiator + Alstra instrument (Biotage). The peptide prepared has a C-terminal carboxylic acid and was synthesized using Fmoc-Gly-Wang resin. The resin was swallowed in DMF for 20 min and fluorenylmethyloxycarbonyl protecting group was deprotected using 20% piperidine in DMF. For the coupling step, Fmoc-amino acid (0.5 mmol) was activated using 0.5mmol of *N,N'*-Diisopropylcarbodiimide (DIC) and 0.5 mmol of ethyl cyanohydroxyiminoacetate (oxyma) in DMF. Coupling was allowed to proceed at room temperature for 5 min. After each coupling step, Fmoc group was deprotected and efficiency of deprotection was automatically controlled by UV absorbance. In the final round, an additional deprotecting step was applied, followed by pre-cleavage wash in Dichloromethane. The peptides were cleaved using a cocktail solution consisting of trifluoroacetic acid (TFA), water, and triisopropylsilane (TIS) in the ratio of 5 mL: 0.5 mL: 0.5 mL. The resin was incubated in the cleavage cocktail for 1 hour at room temperature. Following incubation, the peptide-containing solution was precipitated in 50 mL of ice-cold diethyl ether. The resulting precipitate was collected by centrifugation and subsequently dissolved in 0.5% formic acid (HCOOH) in Milli-Q water. The solution was then lyophilized to obtain the purified peptides. The molecular weight of the peptides was determined using mass spectrometry.

#### 2.2 Modification of peptide with DBCO

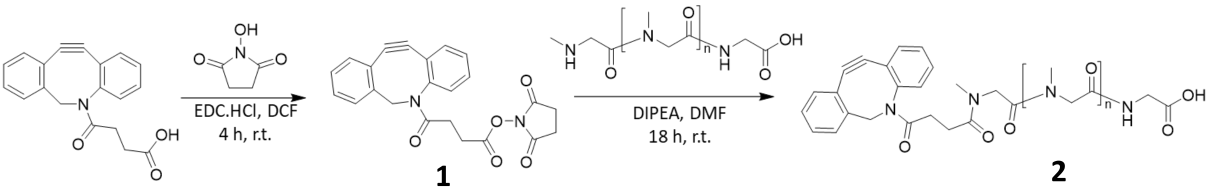


**1** **2**

Synthesis of **1**. 4-(11,12-Didehydrodibenzo[*b,f*]azocin-5(6*H*)-yl)-4-oxobutanoic Acid (20 mg, 0.065 mmol) was dissolved in anhydrous dichloromethane (1 mL) under argon. To this solution, *N*-hydroxysuccinimide (11 mg, 0.098 mmol) and *N*-ethyl-*N'*(3-dimethylaminopropyl)carbodiimide, hydrochloride salt (15 mg, 0.098 mmol) were added and stirred at r.t. for 4 h. The reaction mixture was successively washed with 5% aq. Citric acid (2× 1 mL), 5% sodium hydrogen carbonate solution (2× 1 mL) and brine (1 mL). The organic layer was dried over anhydrous sodium sulfate, filtered and evaporated under reduced pressure. The product **1** was obtained as white solid (26 mg, 100% yield). $^{1}H$ NMR (400 MHz, $CDCl_3$) δ 7.69 (d, 1H), 7.44- 7.25 (m, 7H), 5.18 (d, 2H), 3.68 (d, 1H), 3.02-2.60 (m, 7H), 2.12- 2.04 (m, 1H). $^{13}C$ NMR (400 MHz, $CDCl_3$) δ 170.26, 168.83, 168.30, 151.06, 147.82, 132.30, 129.11, 128.64, 128.38, 127.85, 127.25, 125.56, 123.06, 122.77, 115.05, 107.56, 77.03, 55.60, 29.22, 26.46, 25.54. ESI-MS m/z calculated for $C_{23}H_{18}N_2O_5$ $[M+H]^+$ was 403.1294 and observed 403.1299.

Synthesis of **2.** To a solution of $GSar_n$ peptides (1eq.) in anhydrous *N,N'*-dimethylformamide (200 µL), compound **1** (1.2eq.) and *N,N*-diisopropylethylamine (10eq.) dissolved in 100 µL of anhydrous *N,N'*-dimethylformamide were added. After stirring for 18 h, the reaction mixture was quenched with 700 µL of water. The reaction mixture was evaporated under reduced pressure and re-dissolved in mixture of water and acetonitrile. The product was purified by reverse-phase puriflash column (combiflash system, conditions: A gradient flow of solution A (water + 0.01% TFA) and Solution B (Acetonitrile + 0.01% TFA) was used. For run of 35 min; 0-5 min 100% A; 5-30 min gradient of solution A to 0%, 30-32 min 0% A and 32-35 min 100% A). Fractions were collected and lyophilized and analysed with mass analysis. ESI-MS m/z calculated for $C_{36}H_{43}N_7O_9$ $[M+H]^+$ was 718.3201 and observed 718.3177; $C_{54}H_{73}N_{13}O_{15}Na$ $[M+Na]^+$ was 1166.5247 and observed 1166.5216; $C_{78}H_{113}N_{21}O_{23}$ $[M+H]^+$ was 1712.8396 and observed 1712.8360.

### 2.3 Modification of peptide with chloroalkane ligand

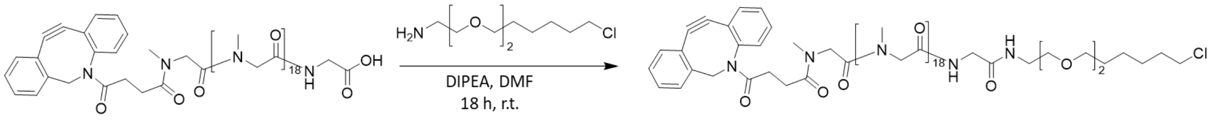


DBCO-GSar19-COOH peptide (2 mg, 1.2 µmol) was dissolved in 100 µL anhydrous *N,N'*-dimethylformamide. To it, pyBOP (9.2 mg, 1.7 µmol) and *N,N*-diisopropylethylamine (0.8 µL, 4.8 µmol) were added and reaction mixture was stirred for 15 min at 4 °C. A solution of chloroalkane ligand amine[a] (0.5 mg, 2.4 µmol) dissolved in DMF (100 µL) was added to reaction mixture. The reaction was stirred for 18 h at r.t. After stirring for 18 h, the reaction mixture was quenched with 250 µL of water.

The reaction mixture was evaporated under reduced pressure and re-dissolved in mixture of water and acetonitrile. The product was purified by reverse phase semi-preparative HPLC column (conditions: A gradient flow of solution A (water + 0.05% TFA) and Solution B (90/10 Acetonitrile/ water + 0.05% TFA) was used. For a run of 35 min; 0-5 min 80% A; 5-30 min gradient of solution A to 0%, 30-32 min 0% A and 32-35 min 80% A; flow rate: 2.5 mL/min). Fractions were collected and lyophilized and analysed with mass analysis. ESI-MS m/z calculated for $C_{36}H_{43}N_7O_9$ $[M+H]^+$ was 1939.9449 and observed 1940.9418.

[a]Chloroalkane ligand amine has been synthesized using previously reported protocol.[3]

### 2.4 Polymer synthesis

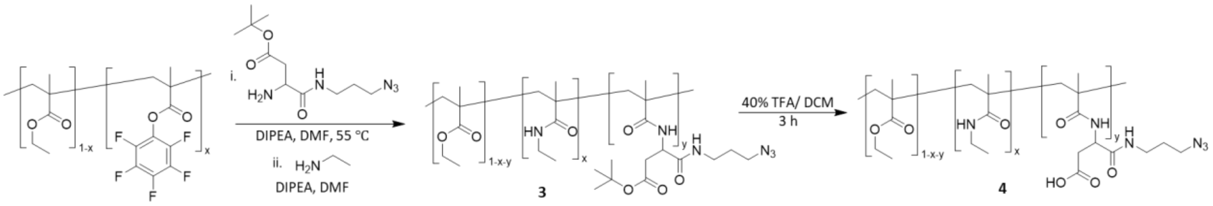


Asp(OtBu)-N3 linker was synthesized as described previously.[4]

Synthesis of **3**. Polymer PEMA-PPF5 (5 eq. of PPF5, 500 mg, 0.2 mol) was dissolved in anhydrous *N,N'*-dimethylformamide (5 mL). To this solution, *N,N*-diisopropylethylamine (30eq., 1.08 mL, 6.2 mol) and Asp(O*t*Bu)-N3 linker (3eq., 278 mg, 0.6 mol) were added. The reaction mixture was stirred for 48 h at 55 °C under argon. The progress of reaction was monitored with $^{19}F$ NMR. The reaction was then quenched with ethylamine (3eq., 0.6 mol, 43 µL) and *N,N*-diisopropylethylamine (4eq., 0.8 mol, 154 µL) dissolved in anhydrous *N,N'*-dimethylformamide (1 mL). The reaction was stirred again for 3 h at r.t. The solvent was then evaporated and was dissolved in minimum of acetonitrile and precipitated in 1:1 MeOH: water and then in methanol. After drying in vacuum the product **3** was obtained as white solid, 160 mg, 32% yield. $^1H$ NMR (400MHz, $CDCl_3$) δ 0.88-1.03 (br d, 3 H), 1.26 (s, 3 H), 1.44 (s, 0.37 H), 1.82-1.92 (br d, 2 H), 4.04 (s, 2 H). (Degree of modification was 56%, calculated from BOC signal in NMR spectra).

Synthesis of **4**. Polymer PEMA-Asp(O*t*Bu)-N3 (114 mg) was dissolved in anhydrous dichloromethane (2.5 mL) and trifluoroacetic acid (1.5 mL) was added at 0 °C. The reaction mixture was warmed to r.t and stirred for 3 h. The solvent was evaporated under reduced pressure and was dissolved in minimum of acetonitrile and precipitated twice in methanol. After drying under vacuum, white solid obtained, 100 mg, 88% yield. $^1H$ NMR (400MHz, $CDCl_3$) δ 0.88-1.03 (br d, 3 H), 1.26 (s, 3 H), 1.44 (s, 0.27 H), 1.82-1.92 (br d, 2 H), 4.04 (s, 2 H).

2.5 HPLC data

1. HPLC trace for GSa5 peptide

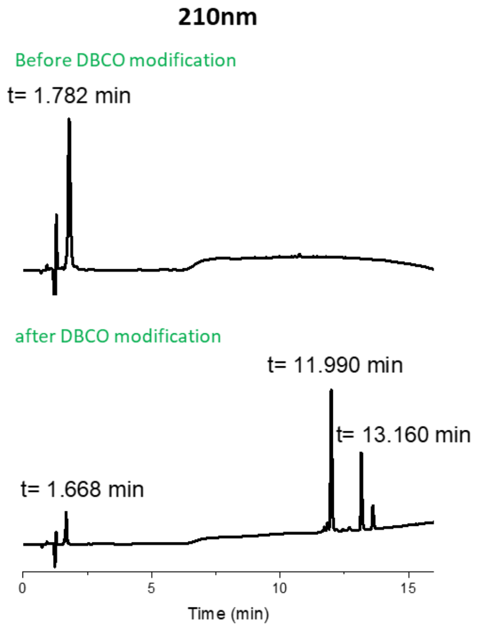


2. HPLC trace for GSar11 peptide

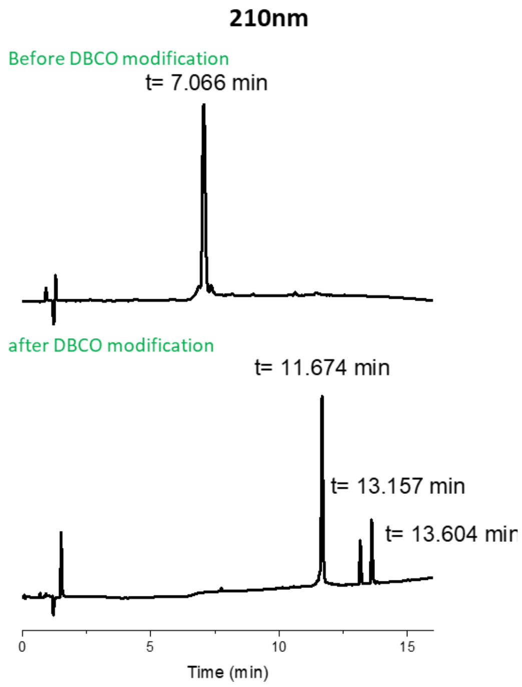

3. HPLC trace for Gsar19 peptide

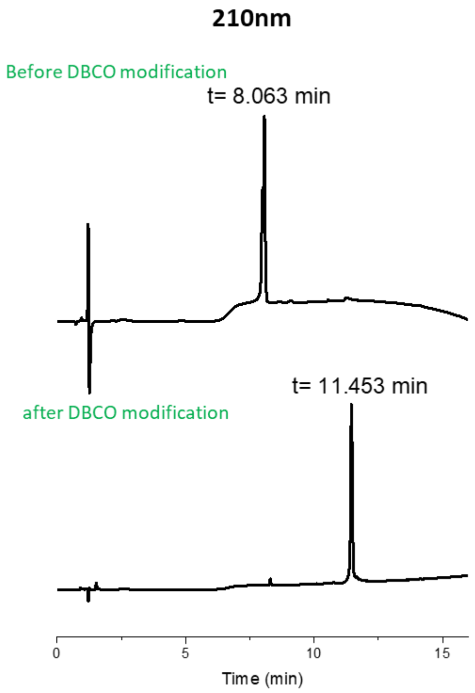

### 2.6 NMR and Mass spectra

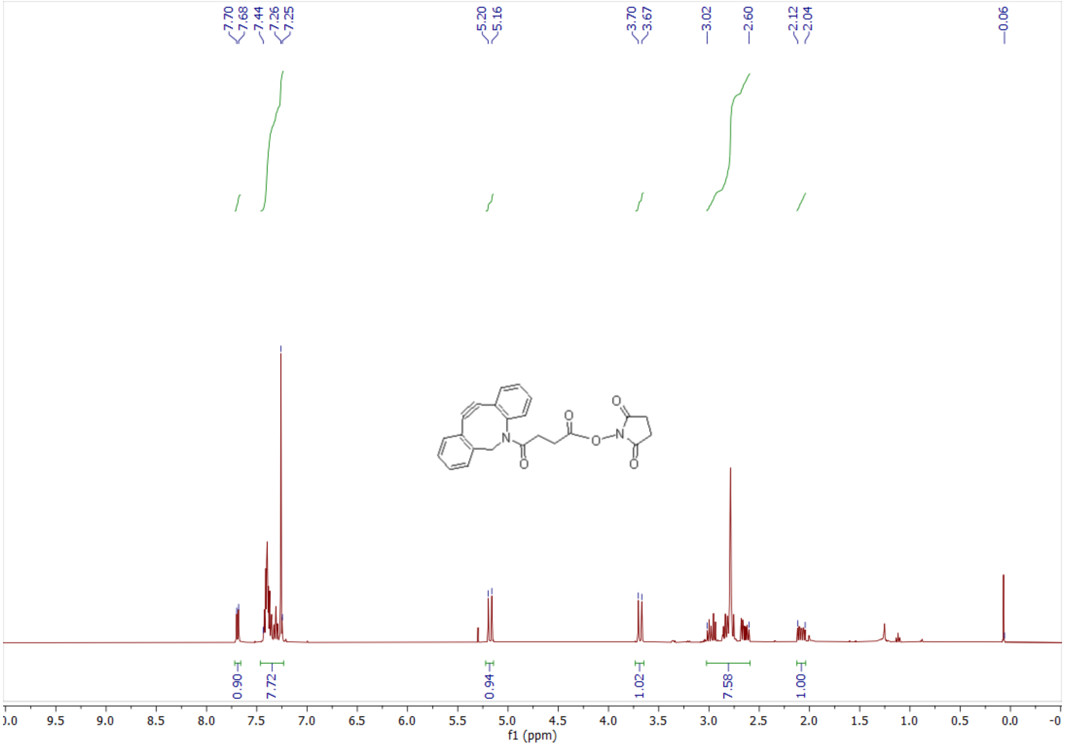


[1]H NMR spectra of DBCO-NHS ester

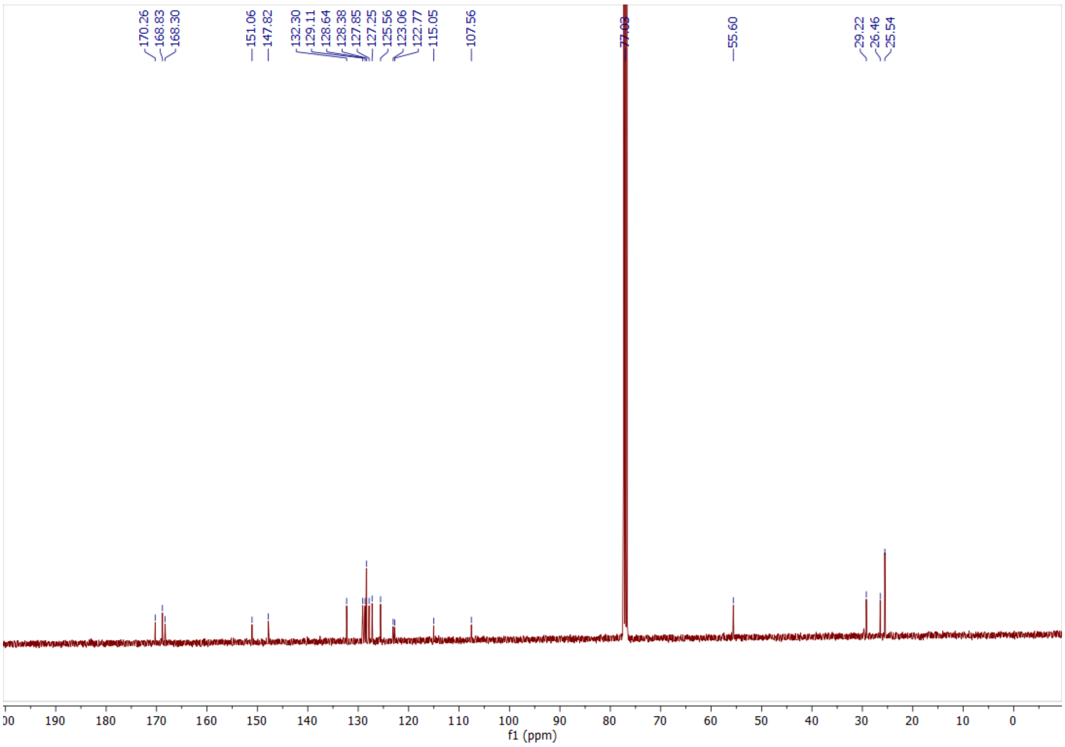


[13]C NMR spectra of DBCO-NHS ester

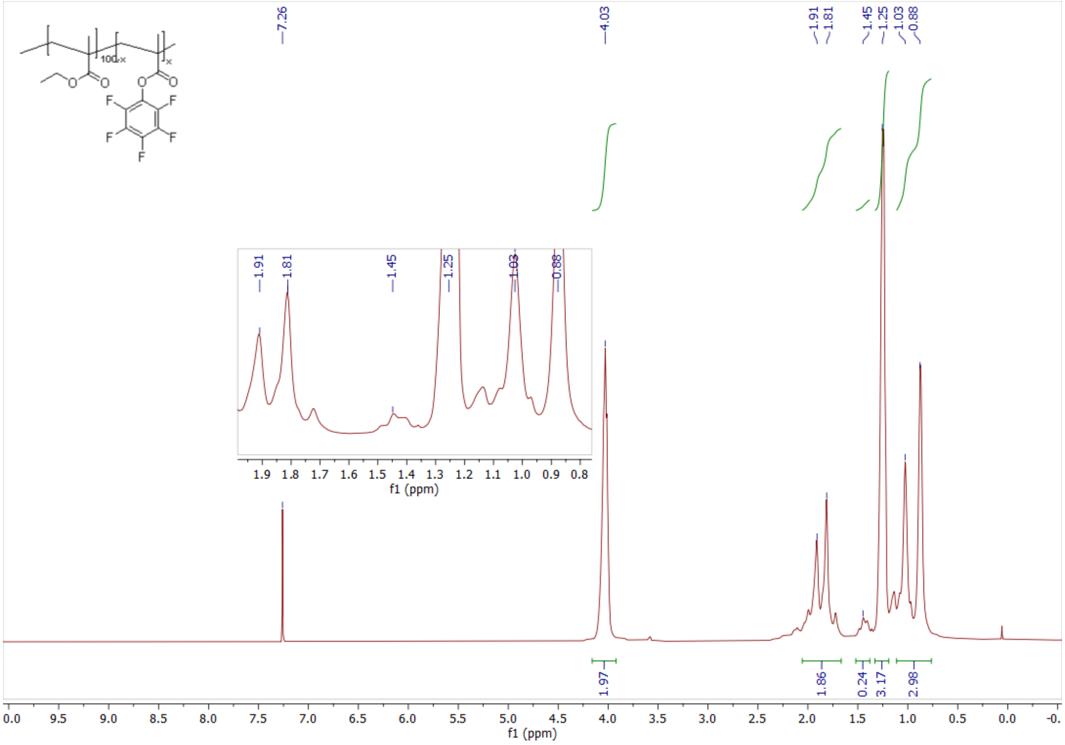


[1]H NMR spectra of PEMA-PPF5 polymer

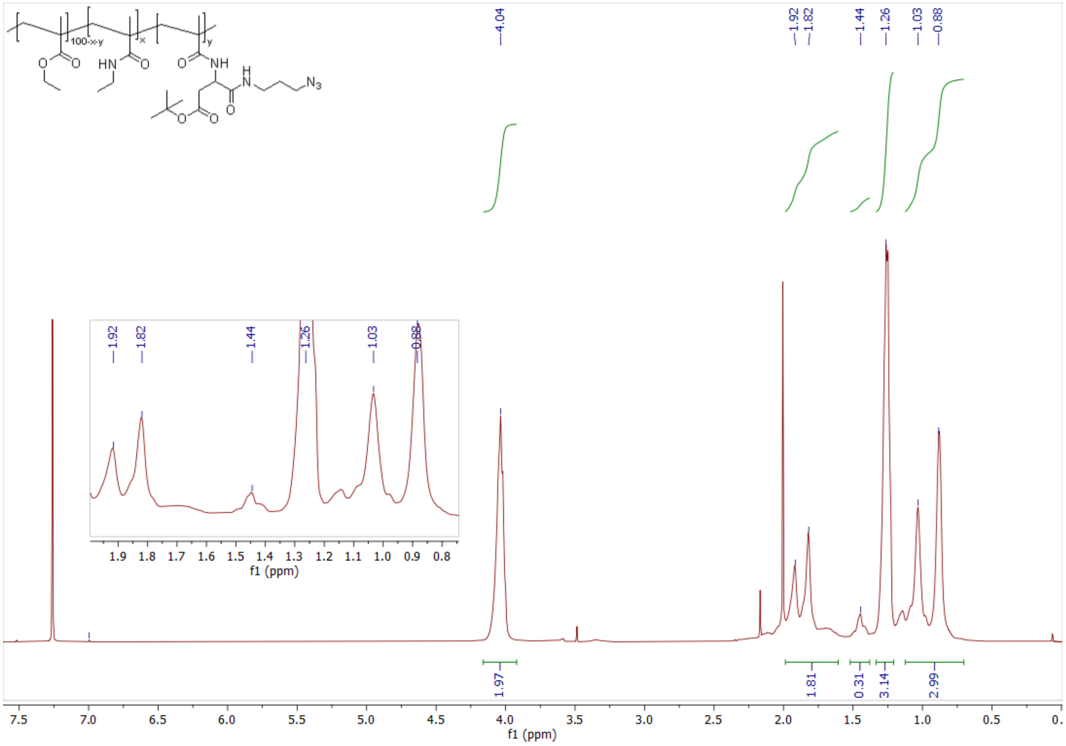


[1]H NMR spectra of PEMA-Asp(OtBu)-N3 polymer (**3**)

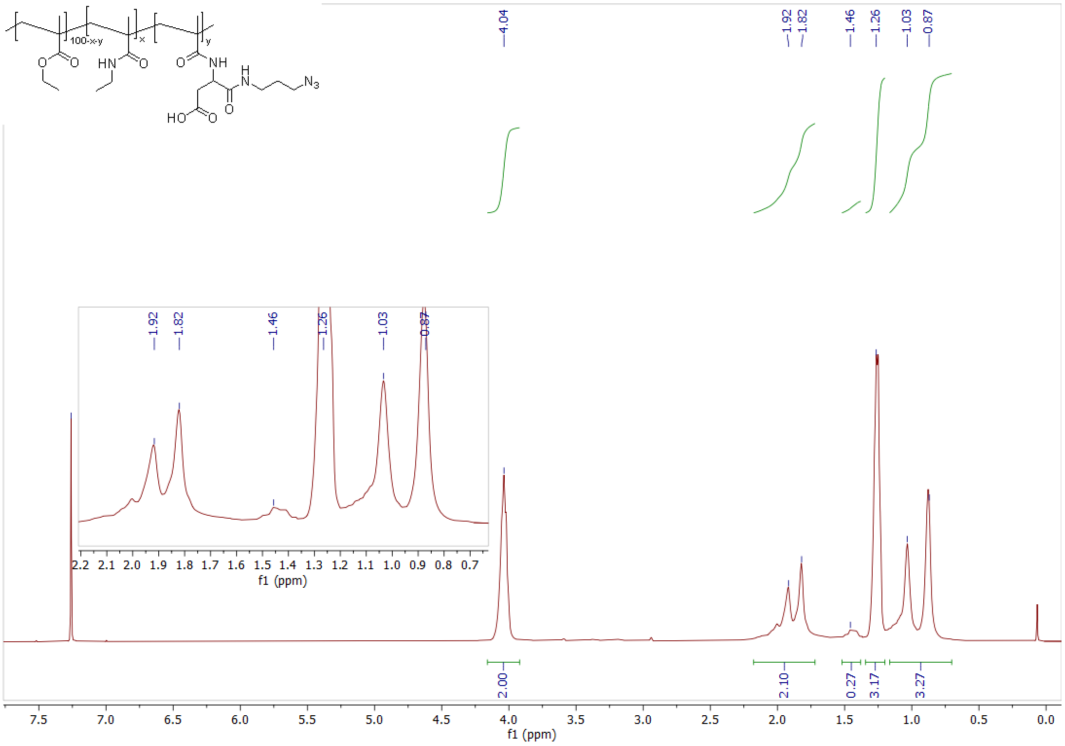


[1]H NMR spectra of PEMA-Asp-N3 polymer (**4**)

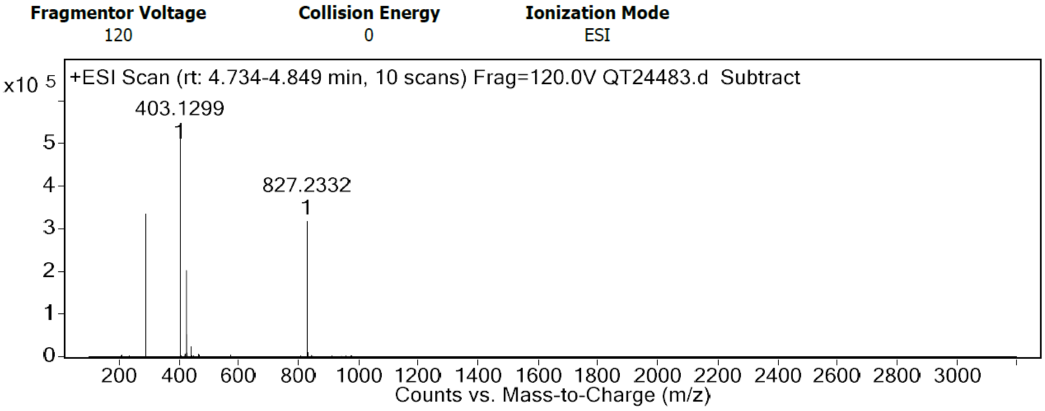


Mass spectra of DBCO-NHS ester (**1**)

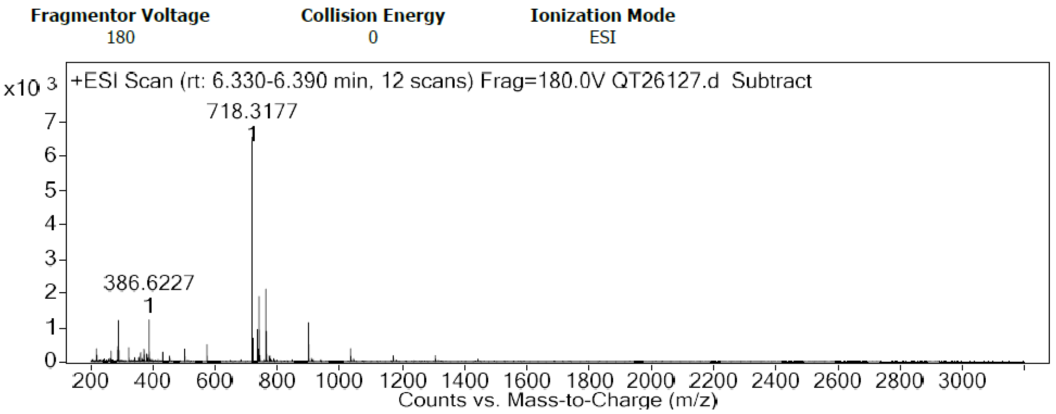


Mass spectra of DBCO-GSar5-COOH peptide

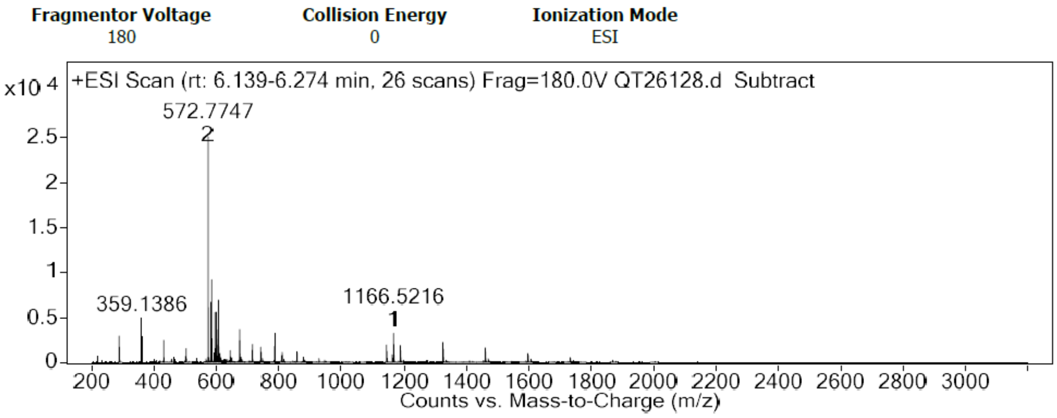


Mass spectra of DBCO-GSar11-COOH peptide

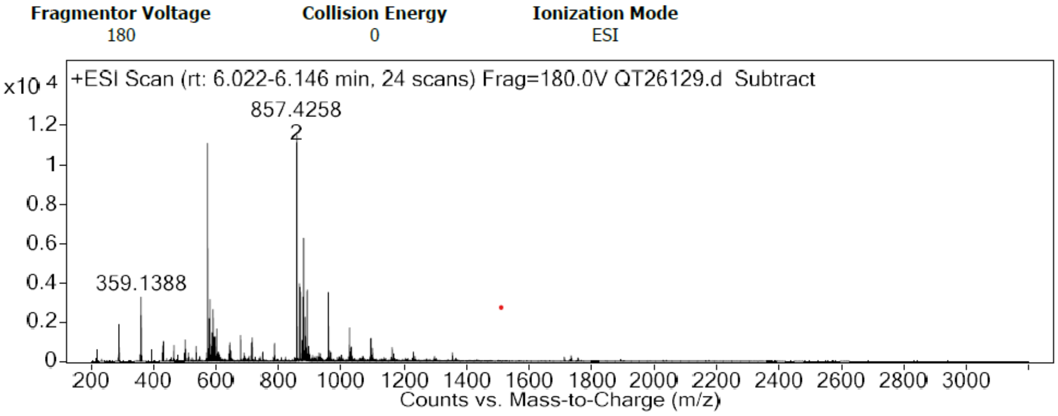


Mass spectra of DBCO-GSar19-COOH peptide

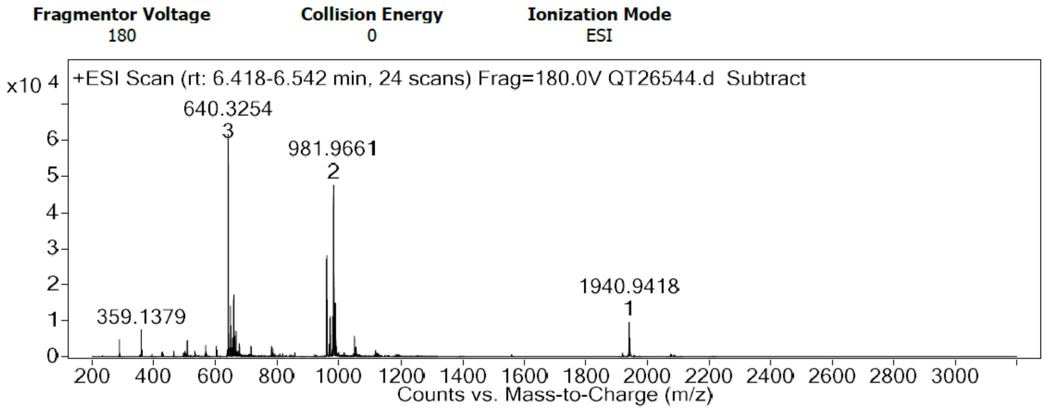


Mass spectra of DBCO-GSar19-Cl peptide

### 2.7 Preparation of dye-loaded Nanoparticles

Polymer PEMA-AspN$_3$ was synthesized previously in the lab. 50 µL of PEMA-AspN$_3$ polymer solution in acetonitrile (2 mg/mL containing 9 wt% R18/F5-TPB dye relative to polymer) was added quickly to 450 µL of 20 mM PB Buffer, pH 7.4 at 21 °C under shaking (thermomixer, 1100 rpm). The particles were then diluted twice.

### 2.8 Synthesis of peptide-functionalized NPs

60 µM of DBCO-GSarn peptide (n = 5, 11, 19) was added to 200 µL of nanoparticles (concentration of azide groups were approx. 26 µM). The reaction was mixed and kept overnight at 40 °C in dark. The reaction was then cooled down to room temperature. The excess of non-reacted peptides was purified by ultracentrifugal filters (Amicon, 0.5 mL, 100kDa) on 10k rpm for 5 min. This procedure was repeated thrice.

### 2.9 Synthesis of Chloride ligand stealth peptide nanoparticle (NPs-GSar19-Cl)

For Synthesis of chloride ligand containing stealth peptide-functionalized NPs; two different ratios of chloride ligand were immobilised on NPs (1% and 10%). For 1% chloride ligand the DBCO-peptides were mixed in ratios 0.6 µM of DBCO-GSar19-Cl and 59.4 µM of DBCO-Gsar19-OH and for 10% chloride ligand; the peptides were mixed in ratio 6 µM of DBCO-GSar19-Cl and 54 µM of DBCO-Gsar19-OH. The reaction was mixed and kept overnight at 40 °C in dark. The reaction was then cooled down to room temperature. The excess of non-reacted peptides was purified by ultracentrifugal filters (Amicon, 0.5 mL, 100kDa) on 10k rpm for 5 min. This procedure was repeated thrice.

### 2.10 Immobilisation of HaloTag GFP protein on NPs-GSar19-Cl

A solution of 10 µM of HaloTag GFP protein was added to 100 µL NPs-GSar19-Cl. The reaction mixture was mixed and incubated for 1 h at 25 °C in the dark. Following incubation, the reaction mixture was diluted five-fold, and excess of unreacted HaloTag GFP was purified using ÄKTA size exclusion chromatography. Aliquots of 500 µL were collected in Eppendorf tubes, and the absorbance and fluorescence spectra of each aliquot were measured. Fluorescence and absorbance were detected in aliquots T11, T12, and T13, with T12 containing the purified sample with immobilised HaloTag-GFP on NPs-GSar19-Cl. The size of NPs was determined using FCS.

### 2.11 Molecular Biology

The plasmid pAG136 for bacterial expression of EGFP was previously described.[5] The plasmid pAG1392 for bacterial expression of Halotag fused to emGFP was constructed by inserting the sequence coding for HaloTag-emGFP in the pET28a vector. Synthetic oligonucleotides used for cloning were purchased from Integrated DNA Technology. PCR reactions were performed with Q5 polymerase (New England Biolabs) in the buffer provided. PCR products were purified using QIAquick PCR purification kit (Qiagen). PCR products and the pET28 plasmid backbone were digested using the enzymes BamH1 and Not1 following the manufacturer protocol (New England Biolabs). The ligation was performed using T4 ligase (New England Biolabs) in the buffer provided. Ligation products were transformed in DH10 E. coli. Small-scale isolation of plasmid DNA was done using QIAprep miniprep kit (Qiagen) from 4 mL of overnight culture supplemented with appropriate antibiotics. Large-scale isolation of plasmid DNA was done using the QIAprep maxiprep kit (Qiagen) from 150 mL of overnight culture supplemented with appropriate antibiotics. Plasmid sequences were confirmed by Sanger sequencing with appropriate sequencing primers (GATC Biotech).

### 2.12 Protein Expression and Purification

*Production of HaloTag-emGFP (resp. EGFP).* Plasmid for the expression of Halotag-emGFP (resp. EGFP) with an N-terminal His-tag under the control of a T7 promoter was transformed in Rosetta *Escherichia coli* competent cells. Bacterial cells were grown at 37 °C in Lysogeny Broth (LB) supplemented with kanamycin (50 µg/mL) until $OD_{600nm}$ = 0.6.  Expression was induced overnight at 16 °C by the addition of isopropyl β-D-1-thiogalactopyranoside (IPTG) (1 mM). Cells were harvested by centrifugation (4000 × g for 20 min at 4 °C) and stored at –30 °C. The cell pellets were resuspended in lysis buffer (PBS supplemented with 2.5 mM $MgCl_2$, protease inhibitor PMSF 1 mM, DNase 0.025 mg/mL) and sonicated (5 min, 20% of amplitude) on ice. The lysate was incubated for 2 h on ice to allow DNA digestion by DNase. Cellular fragments were removed by centrifugation (9,000 × g for 1 h at 4°C). The supernatant was incubated overnight at 4°C by gentle agitation with pre-washed Co-NTA (resp. Ni-NTA) agarose beads in PBS buffer complemented with 10 mM of imidazole. Beads were washed with ten volumes of PBS complemented with 10 mM of imidazole. His-tagged proteins were eluted with five volumes of PBS complemented with 150 mM (respe. 500 mM) of imidazole. The buffer was exchanged to PBS (0.05 M phosphate buffer and 0.150 M NaCl) using PD-10 desalting columns or Midi Trap G-25 (GE Healthcare). The purity of the proteins was evaluated using SDS–PAGE electrophoresis stained with Coomassie blue.

Sequence of 6×His– EGFP:

MGSSHHHHHHSSGLVPRGSHMASVSKGEELFTGVVPILVELDGDVNGHKFSVSGEGEGDATYGKLTLKFICTTGKLP
VPWPTLVTTLTYGVQCFSRYPDHMKQHDFFKSAMPEGYVQERTIFFKDDGNYKTRAEVKFEGDTLVNRIELKGIDFK
EDGNILGHKLEYNYNSHNVYIMADKQKNGIKVNFKIRHNIEDGSVQLADHYQQNTPIGDGPVLLPDNHYLSTQSAL
SKDPNEKRDHMVLLEFVTAAGITLGMDELYK

Sequence of 6×His–HaloTag–emGFP:

MGSSHHHHHHSSGLVPRGSHMASMTGGQQMGRGSMAEIGTGFPFDPHYVEVLGERMHYVDVGPRDGTPVLF
LHGNPTSSYVWRNIIPHVAPTHRCIAPDLIGMGKSDKPDLGYFFDDHVRFMDAFIEALGLEEVVLVIHDWGSALGF
HWAKRNPERVKGIAFMEFIRPIPTWDEWPEFARETFQAFRTTDVGRKLIIDQNVFIEGTLPMGVVRPLTEVEMDHY
REPFLNPVDREPLWRFPNELPIAGEPANIVALVEEYMDWLHQSPVPKLLFWGTPGVLIPPAEAARLAKSLPNCKAVD
IGPGLNLLQEDNPDLIGSEIARWLSTLEISGEPTTEDLYFQSDNAIAVRSASMVSKGEELFTGVVPILVELDGDVNGHK
FSVSGEGEGDATYGKLTLKFICTTGKLPVPWPTLVTTLTYGVQCFARYPDHMKQHDFFKSAMPEGYVQERTIFFKDD
GNYKTRAEVKFEGDTLVNRIELKGIDFKEDGNILGHKLEYNYNSHKVYITADKQKNGIKVNFKTRHNIEDGSVQLADH
YQQNTPIGDGPVLLPDNHYLSTQSALSKDPNEKRDHMVLLEFVTAAGITLGMDELYK

### 3. Supplementary Figures

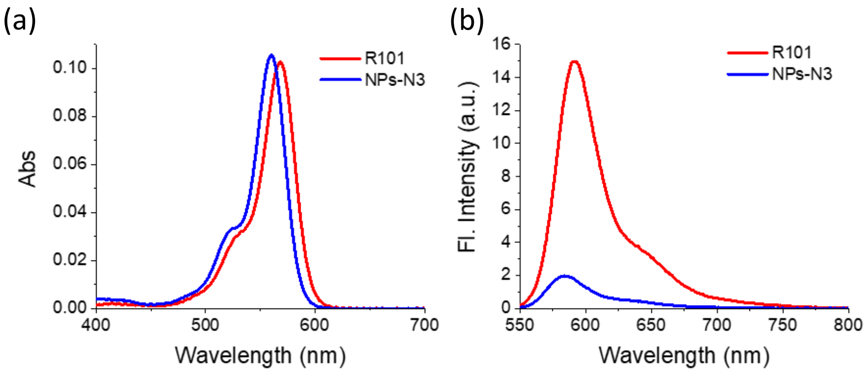


**Figure S1.** Quantum yield. (a) Absorbance spectra (b) Emission spectra for reference (R101 dye) and NPs-N3. Excitation wavelength at 530 nm. Q.Y. of NPs-N3 observed was 0.15.

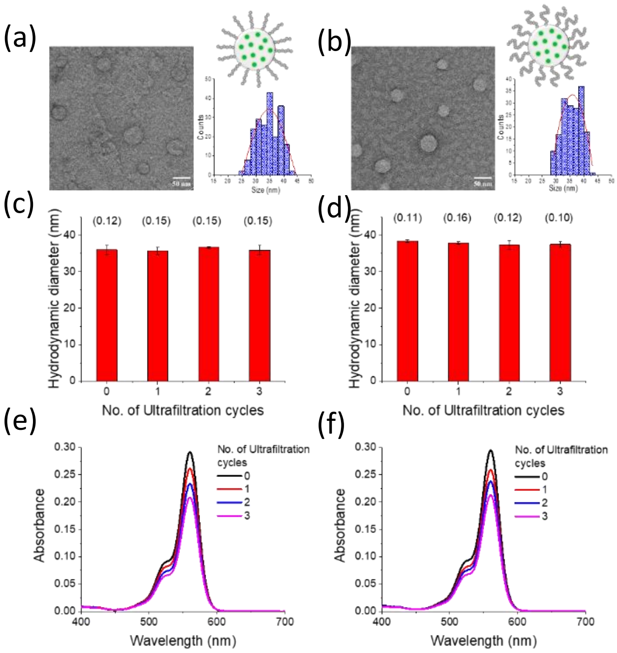


**Figure S2.** Characterisation of Peptide-functionalized NPs. TEM images of (a) NPs-GSar5 and (b) NPs-GSar11 and their corresponding size distribution statistics. Scale bar 50 nm. Size by DLS of NPs after each round of filtration and their polydispersity index (PDI) mentioned in brackets for (c) NPs-GSar5 and (g) NPs-GSar11. The absorbance spectra for (e) NPs-GSar5 (f) NPs-GSar11 after each round of ultrafiltration.

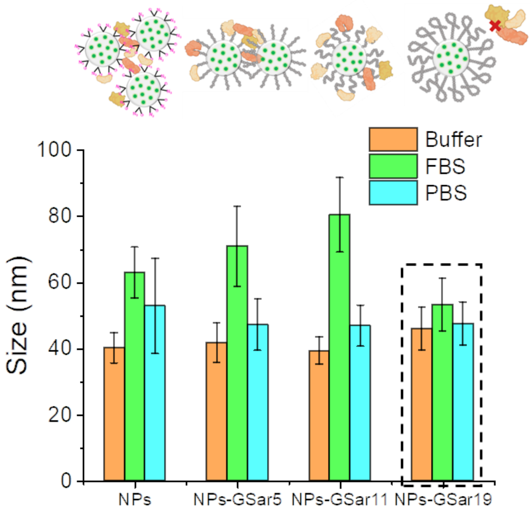


**Figure S3.** Size of NPs (non-modified and modified with 3 different peptides) by Fluorescence Correlation Spectroscopy (FCS) in buffer (orange), in presence of 5% fetal bovine serum (green), and in phosphate buffer saline (Blue).

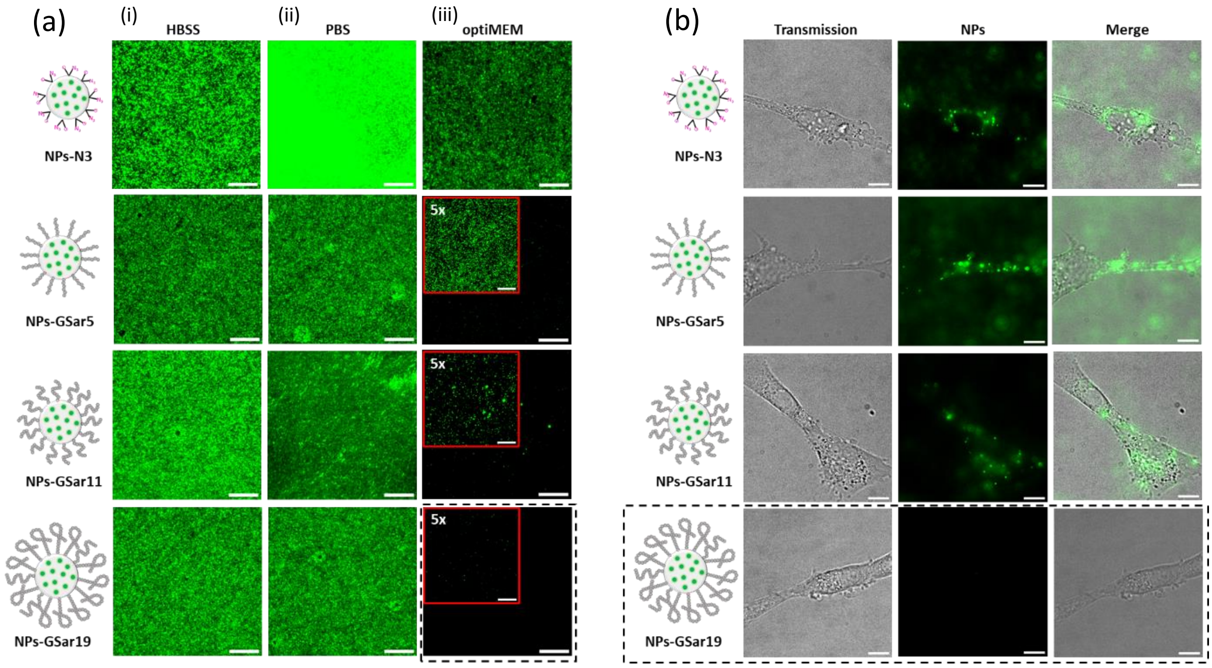


**Figure S4.** (a) Non-specific labelling of NPs-N3 (20 pM) and NPs-GSarn (20 pM) with glass surface using epi-fluorescence microscopy in 3 different biological medias (i) Hank's balanced salt solution (HBSS), (ii) Phosphate buffer saline (PBS) and (iii) optiMEM. The B/C is 1000-6000. Scale bar 10 µm. (b) Non-specific labelling of cell surface of live U87 cells using epi-fluorescence microscopy with NPs-N3 (20pM) and NPs-GSarn (20 pM). The excitation was at 550 nm and integration time was 300 ms. B/C is 20-500. Scale bar 10 µm. The black dotted line shows the best conditions for stealth's NPs.

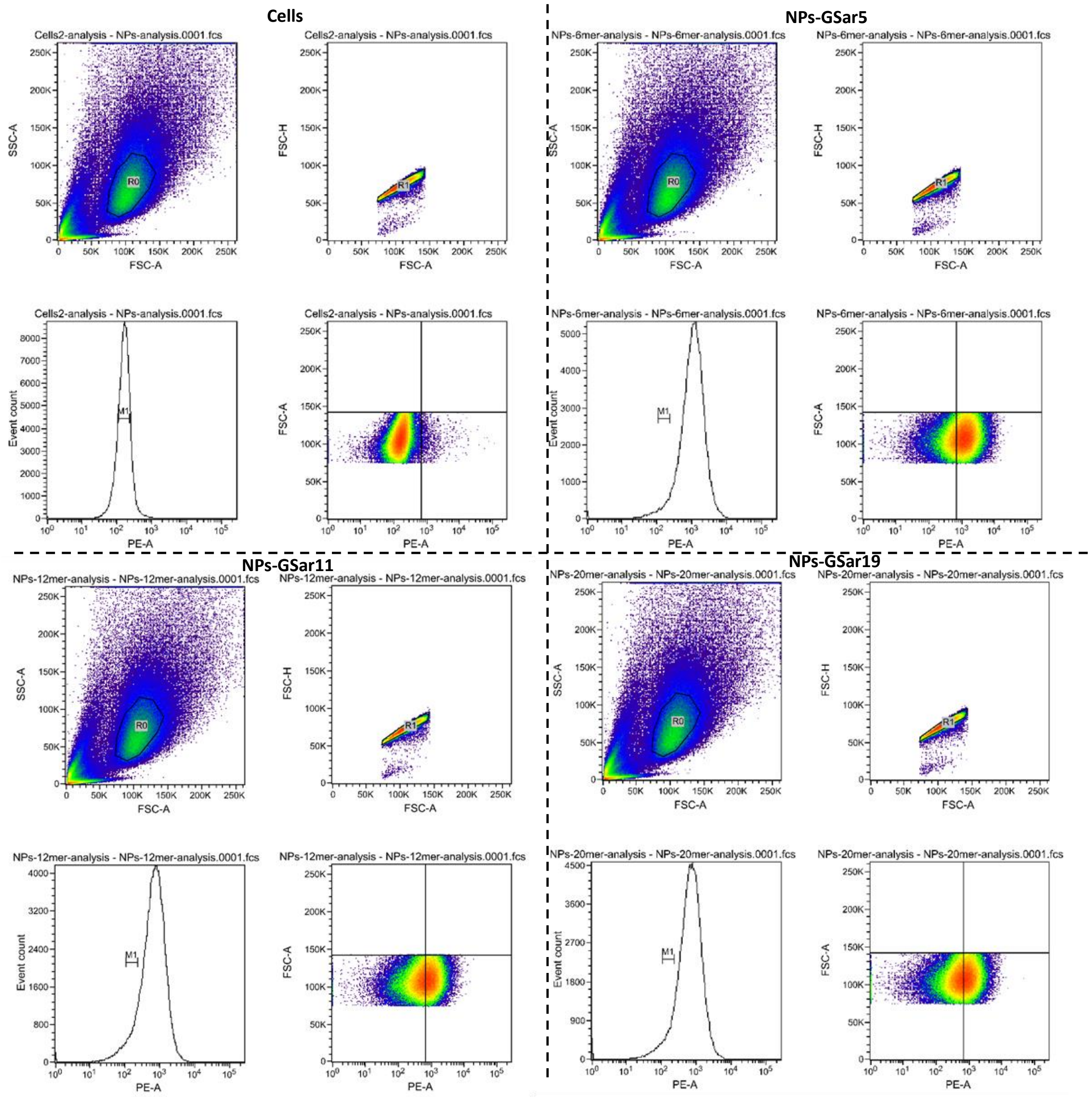


**Figure S5.** Evaluation of nonspecific interactions of NPs with cells by flow cytometry in PBS. Collection gate in side-scatter versus forward scatter dot plot; collection gate in forward-scatter plot versus forward-scatter dot plot; corresponding histogram of gates cells illustrating fluorescence at 550 nm; corresponding dot plot in gated forward-scatter versus fluorescence recorded for (a) Cells (b) NPs-GSar5 (c) NPs-GSar11 (d) NPs-GSar19.

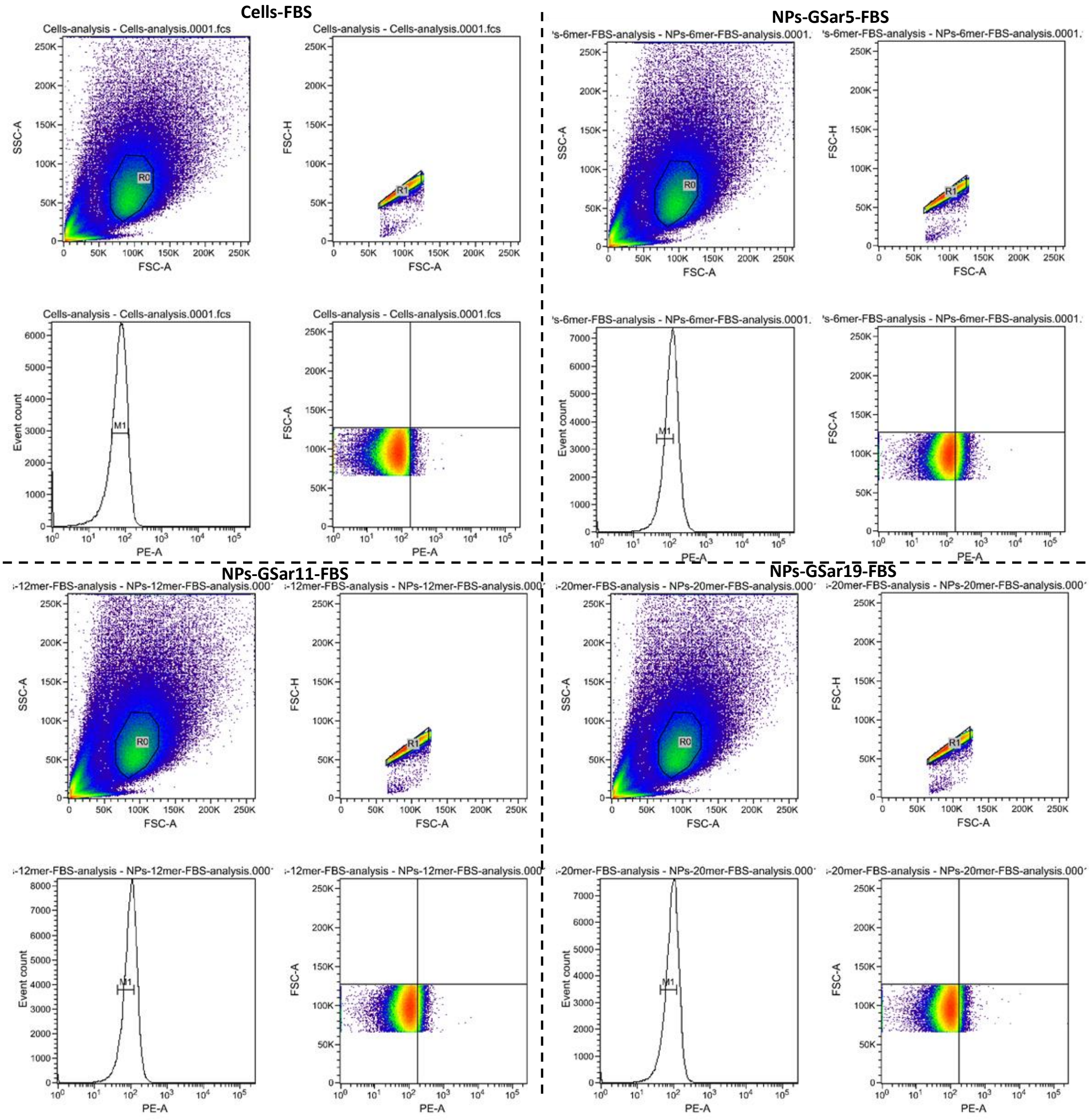


**Figure S6.** Evaluation of nonspecific interactions of NPs with cells by flow cytometry in PBS with 5% FBS. Collection gate in side-scatter versus forward scatter dot plot; collection gate in forward-scatter plot versus forward-scatter dot plot; corresponding histogram of gates cells illustrating fluorescence at 550 nm; corresponding dot plot in gated forward-scatter versus fluorescence recorded for (a) Cells-5% FBS (b) NPs-GSar5-5% FBS (c) NPs-GSar11-5%FBS (d) NPs-GSar19-5%FBS.

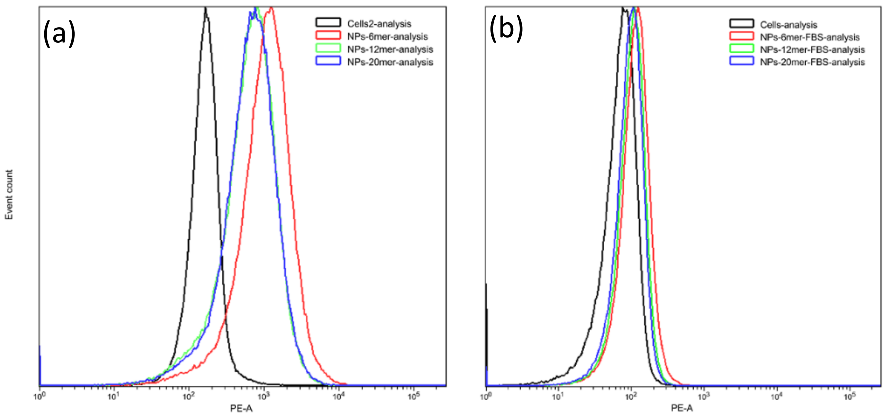


**Figure S7.** Flow cytometry histograms illustrating fluorescence from gated U87 cells (red) with NPs-GSar5 (red); NPs-GSar11 (green); NPs-GSar19 (blue) in (a) absence of FBS (b) presence of FBS.

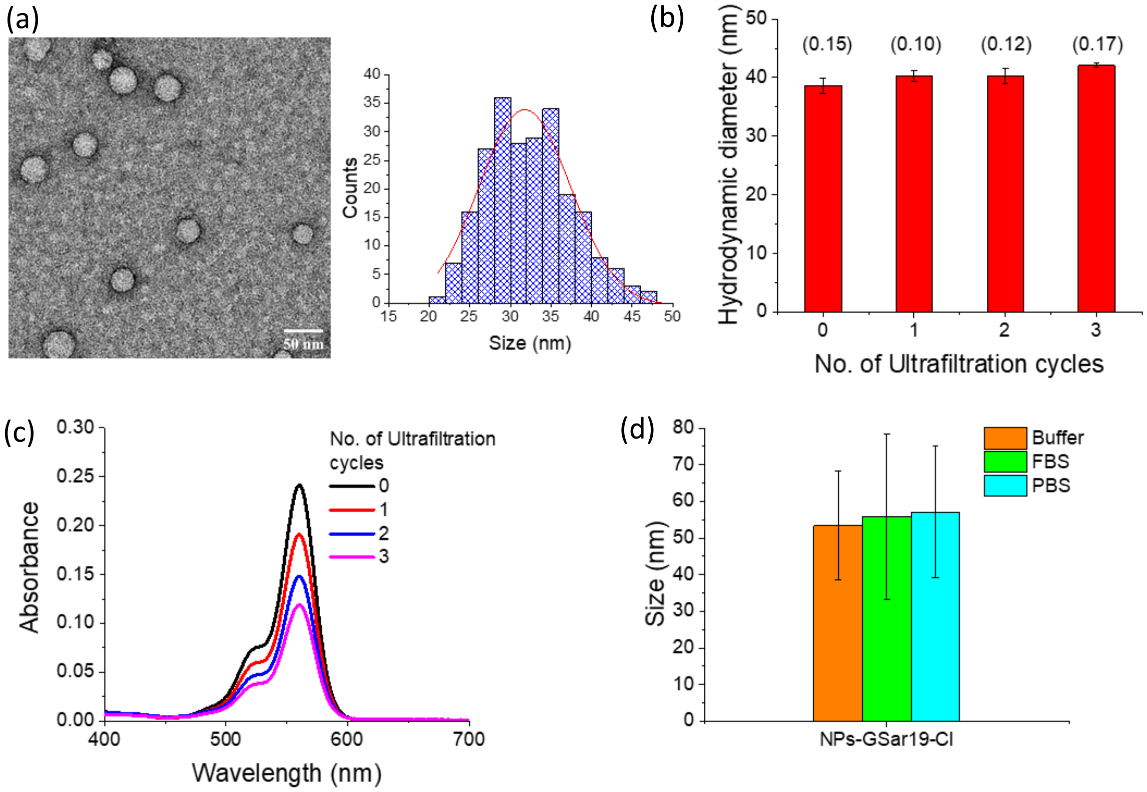


**Figure S8.** (a) TEM images of NPs-GSar19-Cl and its corresponding size distribution statistics. Scale bar 50 nm. (b) Size by DLS with polydispersity index in brackets and (c) absorbance spectra for NPs-GSar19-Cl after each round of ultrafiltration. (d) Size of NPs by FCS in presence of buffer (orange), 5% FBS (green), and PBS (blue).

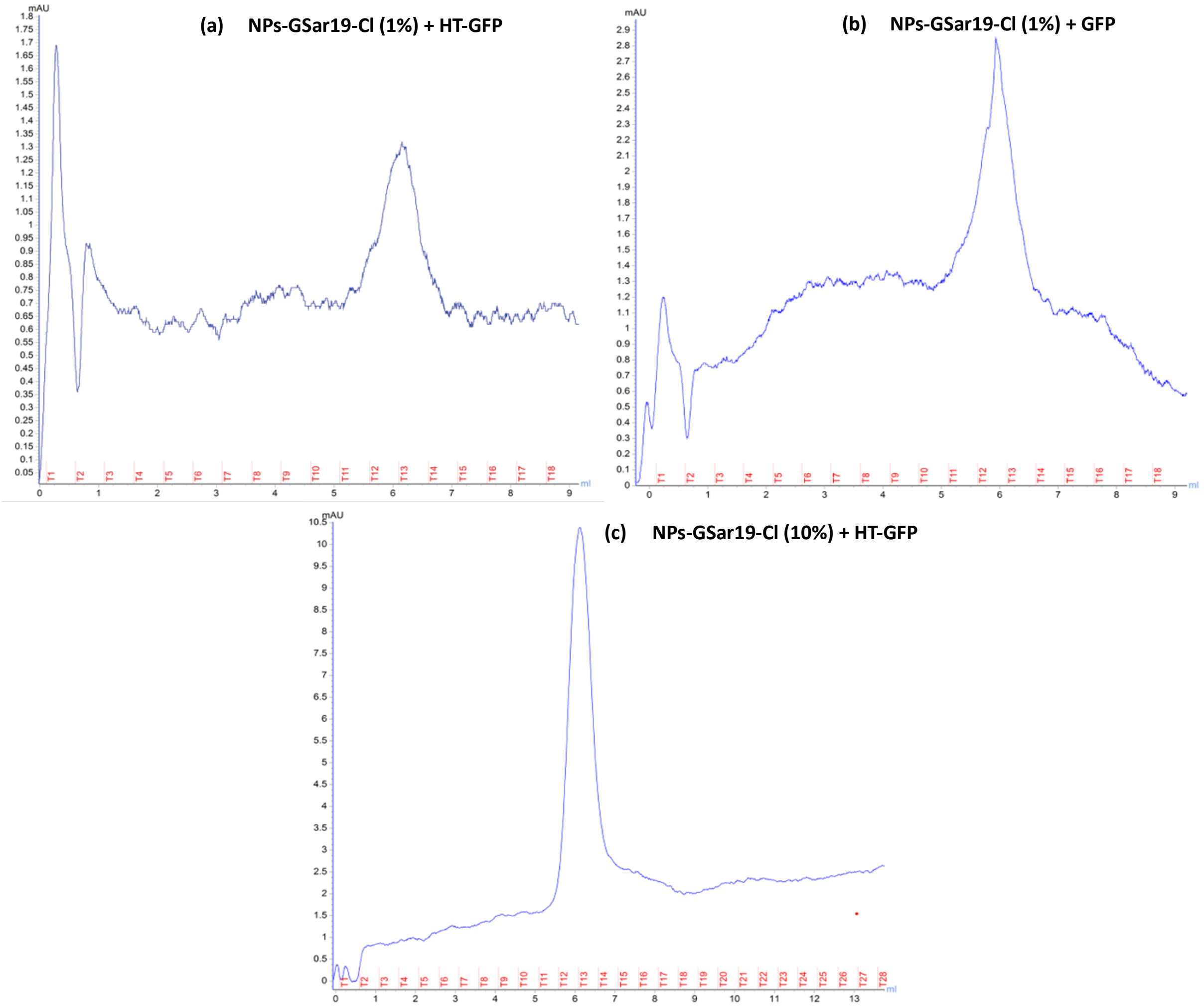


**Figure S9.** UV chromatograms at 280 nm for ÄKTA size exclusion chromatography of (a) NPs-Gar19-Cl(1%) + HT-GFP (b) NPs-Gar19-Cl(1%) + GFP (c) NPs-GSar19-Cl(10%) + HT-GFP.

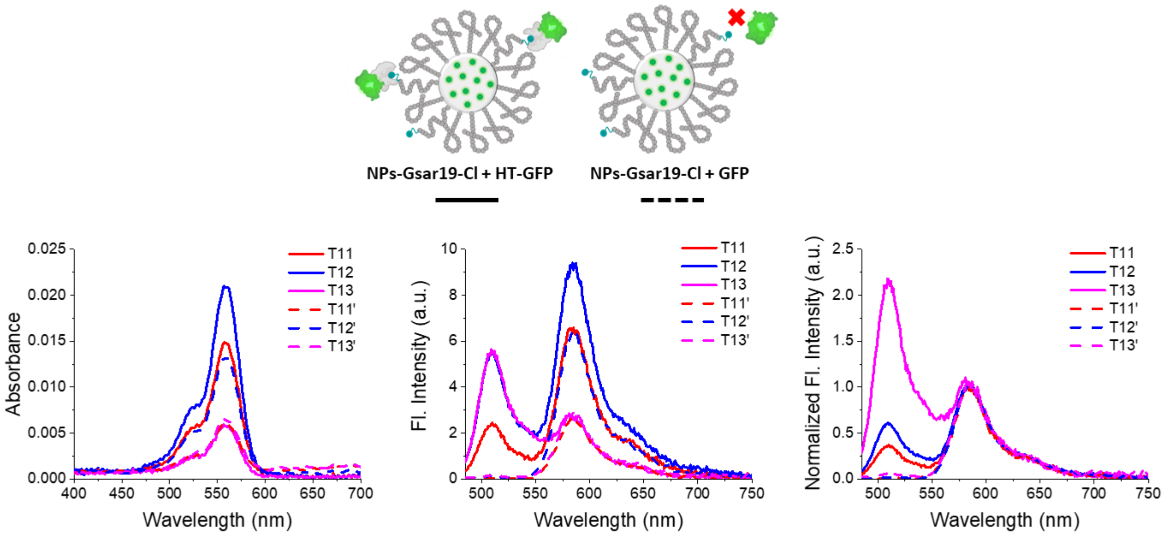


**Figure S10.** NPs-GSar19-Cl (1%) incubated with HaloTag GFP (solid line) and GFP (dashed line) purified by ÄKTA size exclusion chromatography (a) absorbance spectra (b) fluorescence spectra excited at 470 nm (c) normalized fluorescence intensity spectra for the sample collected in tubes 11 (red), 12 (blue), and 13 (magenta).

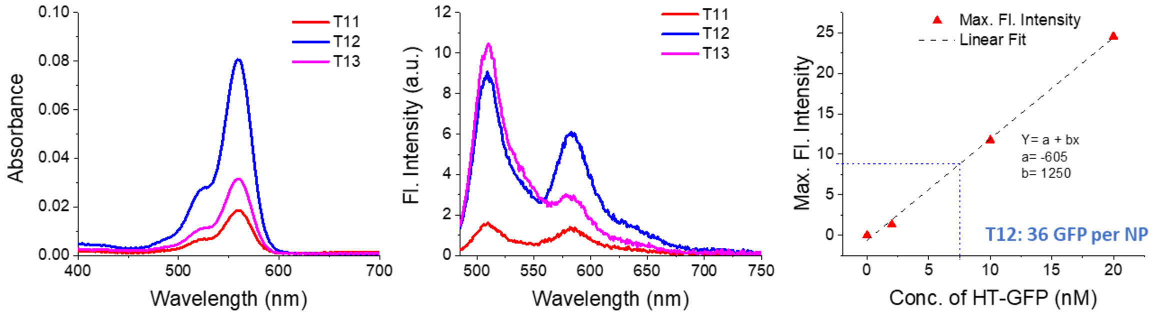


Figure S11. NPs-GSar19-Cl (10%) incubated with HaloTag GFP purified by ÄKTA size exclusion chromatography (a) absorbance spectra (b) fluorescence spectra excited at 470 nm for the sample collected in fractions 11 (red), 12 (blue), and 13 (magenta). (c) Calibration curve for the concentration of HaloTag GFP versus its recorded fluorescence intensity maxima at 510 nm.

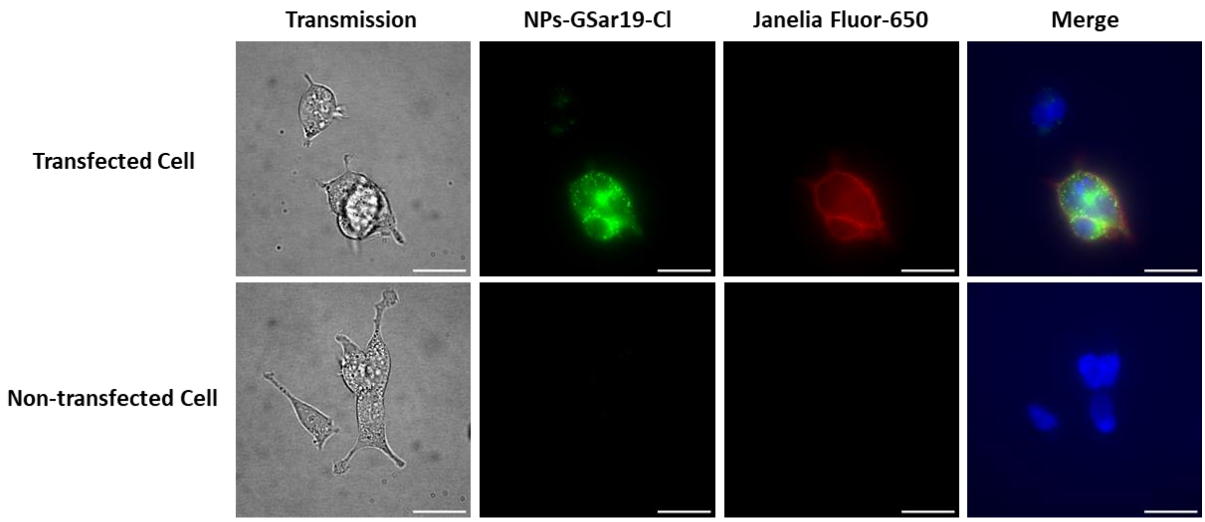


**Figure S12.** Microscope images for labelling of HaloTag proteins with 200 pM NPs-GSar19-Cl in presence of 5% FBS (green colour) and 1 nM commercial Janelia Fluor-650 dye (red colour) expressed on cell surface of transfected HEK293T cells (upper panel) and non-transfected HEK cells (lower panel). The nucleus was labelled with Hoechst dye in blue colour. B/C for green channel 30-650 and 100-5000 for red channel. Scale bar 25 µm.

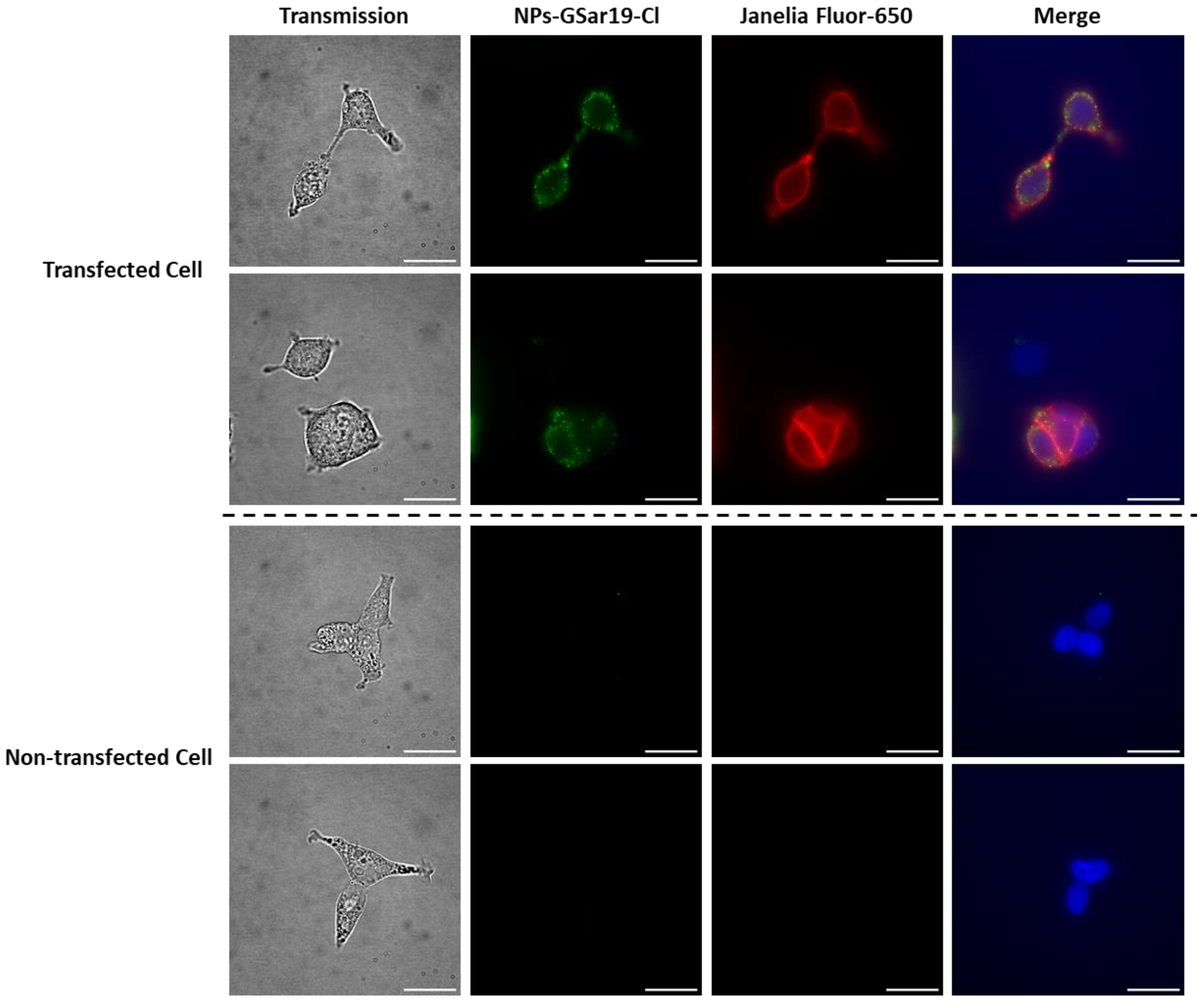


**Figure S13.** Microscope images for labelling of HaloTag proteins with 200 pM NPs-GSar19-Cl in presence of 0.01 g/L Tween 20 (green colour) and 1 nM commercial Janelia Fluor-650 dye (red colour) expressed on cell surface of transfected HEK293T cells (upper panel) and non-transfected HEK cells (lower panel). The nucleus was labelled with Hoechst dye in blue colour. B/C for green channel 30-650 and 100-5000 for red channel. Scale bar 25 µm.

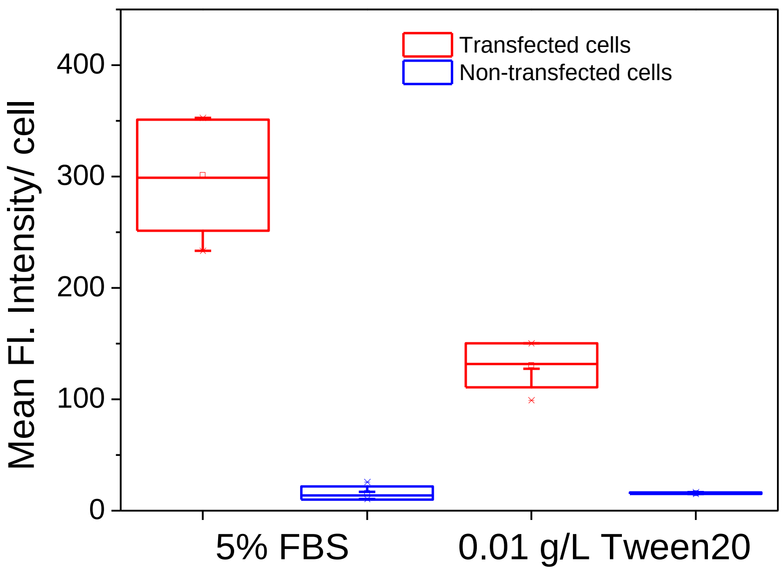


**Figure S14.** Analysis of microscopy images for labelling of HaloTag proteins with 200 pM NPs-GSar19-Cl in transfected cells showing positive signal with Janelia Fluor-650 and control non-transfected cells. Five cells were analysed per condition.

### 4. Tables

**Table S1.** FCS parameters obtained for studied samples in Figure 3 with single-component fit (T+3D model). It is a model that includes triplet and one emissive specie with 3D diffusion; n is the number of emissive species; τ_diff [ms] is diffusion correction time; cpp [kHz] counts per particle; conc. (nM) concentration of emissive species; Size (nm) hydrodynamic diameter calculated from τ_diff; and brightness calculated from cpp.

| Sample | avg. signal [kHz] | n | τ_diff [ms] | cpp [kHz] | Conc. (nM) | Size (nm) | Brightness |
|---|---|---|---|---|---|---|---|
| TMR | 32.66 | 62.16 | 0.04 | 0.53 | 50.00 | 1.00 | 1.00 |
| NPs | 225.88 | 3.46 | 1.28 | 65.08 | 2.78 | 34.20 | 123.63 |
| NPs-FBS | 118.71 | 57.16 | 2.02 | 2.08 | 45.98 | 53.86 | 3.95 |
| NPs-NaCl | 346.91 | 7.57 | 1.42 | 45.97 | 6.09 | 37.76 | 87.34 |
| NPs-GSar5 | 184.60 | 2.78 | 1.34 | 66.49 | 2.24 | 35.63 | 126.31 |
| NPs-GSar5-FBS | 193.33 | 1.62 | 1.82 | 118.64 | 1.30 | 48.51 | 225.38 |
| NPs-GSar5-NaCl | 141.89 | 1.97 | 1.37 | 70.26 | 1.58 | 36.17 | 133.47 |
| NPs-GSar11 | 110.77 | 1.69 | 1.32 | 65.86 | 1.36 | 35.30 | 125.12 |
| NPs-GSar11-FBS | 96.45 | 4.25 | 1.47 | 22.63 | 3.42 | 39.12 | 42.99 |
| NPs-GSar11-NaCl | 92.88 | 1.46 | 1.31 | 62.90 | 1.17 | 34.58 | 119.49 |
| NPs-GSar19 | 138.85 | 2.16 | 1.47 | 63.22 | 1.73 | 39.07 | 120.10 |
| NPs-GSar19-FBS | 155.10 | 1.66 | 1.45 | 94.95 | 1.34 | 38.43 | 180.39 |
| NPs-GSar19-NaCl | 98.97 | 1.61 | 1.38 | 60.52 | 1.29 | 36.77 | 114.98 |

**Table S2.** FCS parameters obtained for studied samples in Figure S3 with single-component fit (T+3D model). It is a model that includes triplet and one emissive specie with 3D diffusion; n is the number of emissive species; τ_diff [ms] is diffusion correction time; cpp [kHz] counts per particle; conc. (nM) concentration of emissive species; Size (nm) hydrodynamic diameter calculated from τ_diff; and brightness calculated from cpp.

| Sample | avg. signal [kHz] | n | τ_diff [ms] | cpp [kHz] | Conc. (nM) | Size (nm) | Brightness |
|---|---|---|---|---|---|---|---|
| TMR | 55.69 | 19.72 | 0.04 | 2.81 | 50.00 | 1.00 | 1.00 |
| TMR-2 (NaCl) | 102.69 | 21.87 | 0.04 | 4.70 | 50.00 | 1.00 | 1.00 |
| NPs | 160.72 | 1.08 | 1.50 | 149.25 | 2.73 | 40.09 | 53.04 |
| NPs-FBS | 127.75 | 1.12 | 2.35 | 115.51 | 2.84 | 62.87 | 41.05 |
| NPs-NaCl | 46.92 | 0.19 | 2.20 | 251.63 | 0.43 | 52.79 | 53.57 |
| NPs-GSar5 | 98.79 | 0.60 | 1.56 | 164.09 | 1.53 | 41.69 | 58.31 |
| NPs-GSar5-FBS | 98.62 | 0.96 | 2.65 | 104.16 | 2.44 | 70.86 | 37.01 |
| NPs-GSar5-NaCl | 44.23 | 0.42 | 1.71 | 101.24 | 0.95 | 47.11 | 21.55 |
| NPs-GSar11 | 105.39 | 0.74 | 1.47 | 144.85 | 1.87 | 39.28 | 51.47 |
| NPs-GSar11-FBS | 82.23 | 2.38 | 3.01 | 34.20 | 6.03 | 80.36 | 12.15 |
| NPs-GSar11-NaCl | 95.13 | 0.59 | 1.70 | 158.42 | 1.36 | 46.80 | 33.73 |
| NPs-GSar19 | 130.22 | 0.88 | 1.71 | 151.56 | 2.23 | 45.86 | 53.86 |
| NPs-GSar19-FBS | 131.94 | 1.29 | 1.99 | 103.67 | 3.27 | 53.26 | 36.84 |
| NPs-GSar19-NaCl | 54.76 | 0.43 | 1.71 | 124.93 | 0.98 | 47.46 | 26.60 |

**Table S3.** FCS parameters obtained for studied samples in Figure S8 with single-component fit (T+3D model).[a]

| Samples | avg. signal [kHz] | n | τ_diff [ms] | cpp [kHz] | Conc. (nM) | Size (nm) | Brightness |
|---|---|---|---|---|---|---|---|
| TMR | 56.17 | 45.34 | 0.04 | 1.25 | 50.00 | 1.00 | 1.00 |
| NPs-GSar19-Cl | 21.19 | 0.29 | 2.41 | 73.12 | 0.32 | 53.37 | 58.39 |
| NPs-GSar19-Cl-FBS | 22.20 | 0.36 | 2.38 | 59.53 | 0.40 | 55.81 | 47.53 |
| NPs-GSar19-Cl-NaCl | 17.06 | 0.23 | 2.43 | 74.34 | 0.25 | 57.14 | 59.36 |

[a]It is a model that includes triplet and one emissive specie with 3D diffusion; n is the number of emissive species; τ_diff [ms] is diffusion correction time; cpp [kHz] counts per particle; conc. (nM) concentration of emissive species; Size (nm) hydrodynamic diameter calculated from τ_diff; and brightness calculated from cpp.

**Table S4.** FCS parameters obtained for studied samples in Figure 5 with single-component fit (T+3D model).[a]

| Samples | avg. signal [kHz] | n | τ_diff [ms] | cpp [kHz] | Conc. (nM) | Size (nm) | Brightness |
|---|---|---|---|---|---|---|---|
| TMR08 | 80.62 | 32.10 | 0.04 | 2.50 | 50.00 | 1.00 | 1.00 |
| NPs | 117.31 | 2.43 | 1.38 | 48.49 | 3.78 | 32.67 | 19.37 |
| NPs+ HT-GFP | 7.34 | 0.23 | 1.87 | 30.57 | 0.36 | 44.21 | 12.21 |
| NPs+ GFP | 3.33 | 0.10 | 1.55 | 34.33 | 0.16 | 36.72 | 13.71 |

[a] It is a model that includes triplet and one emissive specie with 3D diffusion; n is the number of emissive species; τ_diff [ms] is diffusion correction time; cpp [kHz] counts per particle; conc. (nM) concentration of emissive species; Size (nm) hydrodynamic diameter calculated from τ_diff; and brightness calculated from cpp.
